# The global proteome of *Streptococcus pneumoniae* EF3030 under nutrient-defined *in vitro* conditions

**DOI:** 10.1101/2025.04.07.647550

**Authors:** Supradipta De, Larissa M. Busch, Gerhard Burchhardt, Manuela Gesell Salazar, Rabea Schlüter, Leif Steil, Uwe Völker, Sven Hammerschmidt

**Affiliations:** Department of Molecular Genetics and Infection Biology, Interfaculty Institute of Genetics and Functional Genomics, University of Greifswald, Greifswald, Germany; Department of Functional Genomics, Interfaculty Institute of Genetics and Functional Genomics, University Medicine Greifswald, Germany; Imaging Center of the Department of Biology, University of Greifswald, Greifswald, Germany

**Keywords:** *Streptococcus pneumoniae*, minimal growth medium, scanning electron microscopy, proteome, bacterial morphology

## Abstract

*Streptococcus pneumoniae* is a pathobiont that colonises the upper respiratory tract of humans without causing symptoms but can cause a range of life-threatening diseases including pneumonia, sepsis, and meningitis. It also causes less severe, non-invasive infections such as otitis media and sinusitis. This bacterium thrives in the nasopharynx, where nutrient availability is limited, and has adapted to this environment by developing mechanisms to survive host stress and regulate protein abundance. To study the molecular biology of *S. pneumoniae* under *in vitro* and infection-related conditions, a suitable cultivation medium is essential for reproducible experiments. In this study, we optimized a chemically-defined minimal medium that mimics the *in vivo* nutrient-limited condition and used it for proteome analysis. This optimized medium not only shortened the lag phase but also improved the growth of *S. pneumoniae* clinical isolates and other streptococcal species. We applied this medium to analyse the global proteome of the pneumococcal colonising strain EF3030, focusing on the transition from the early to late log phase. Our proteomic analysis revealed distinct patterns of protein abundance in different functional categories including metabolism, amino acid synthesis, natural competence, RNA synthesis, cell wall synthesis, protein degradation, and stress responses. Notably, choline-binding protein CbpD, competence factors ComGA and ComEA as well as proteins involved in processing internalized single-stranded DNA (ssDNA) such as Dpr and DprA were higher in abundance in the late log phase. This proteomic profiling provides valuable insights into the pathophysiology of strain *S. pneumoniae* EF3030 under defined nutrient conditions.

## 1 Introduction

*Streptococcus pneumoniae,* also known as pneumococcus, exhibits a range of growth behaviours, including planktonic growth as diplococci, in chains, or in biofilms. Biofilm formation is particularly significant in the asymptomatic colonisation of the upper respiratory tract (URT), where *S. pneumoniae* predominantly resides (Chao et al., 2014; Gilley and Orihuela, 2014). However, any alteration in the URT can trigger the dissemination of pneumococci and the onset of pathogenesis (Simell et al., 2012). The ability of pneumococci to cross host tissue barriers is a critical factor in its potential to cause both non-invasive and invasive diseases such as otitis media, sinusitis, pneumonia, septicaemia, or meningitis, posing significant global health challenges (Bogaert et al., 2004; Yao and Yang, 2008; Hammerschmidt, 2006; Cartwright, 2002).

The growth of *S. pneumoniae* is significantly influenced by a diverse array of environmental and nutrient factors that can vary widely between host compartments and controlled growth conditions (Härtel et al., 2012; Schulz et al., 2014; Carvalho et al., 2013; Leonard et al., 2018; Sanchez-Rosario and Johnson, 2021; Burghout et al., 2013; Fatykhova et al., 2024; Schulz and Hammerschmidt, 2013; Härtel et al., 2011). Carbohydrates and amino acids, particularly, play a decisive role in the success of the bacteria in encountering and adapting to different host niches, and in causing infectious diseases (Härtel et al., 2012). Within its primary niche, the URT, the availability of nutrients, including nasal transition metal levels, differs substantially from other environments such as blood stream and lungs, which influences the phenotype and pathophysiology of pneumococci (van Beek et al., 2020; Krismer et al., 2014; De, S., Hakansson, A.P., 2023; Im et al., 2022).

A previous study by our group (Schulz et al., 2014) introduced several supplements to RPMI 1640, including uracil, adenine, glycine, choline chloride, phosphate, carbonate, and glucose, to improve the cultivation of streptococcal strains in a minimal medium. Notably, nasal fluid contains 10-fold higher concentrations of calcium and magnesium compared to iron, copper and zinc (van Beek et al., 2020). These variations in metal availability have a profound impact on *S. pneumoniae* growth and metabolism, for example, the balance between iron and manganese plays a key role in regulating peroxide levels via the Fenton reaction, thereby influencing the oxidative stress response and contributing to pneumococcal pathogenicity (Ong et al., 2013). Understanding host compartment-specific nutrient variations provides valuable insights into bacterial metabolism and the mechanisms that enable *S. pneumoniae* to adapt to different environments, thereby improving *in vitro* models that more accurately mimic the host environment.

In the nasopharynx, *S. pneumoniae* encounters significant challenges in overcoming mucus entrapment and evading the immune defence systems of the host. The capsular polysaccharide (CPS) helps pneumococci to bypass the electrostatic repulsion from mucus, facilitating bacterial adhesion to epithelial cells (Nelson et al., 2007). However, the capsule can also conceal key surface proteins involved in recognising host cells. To overcome this, *S. pneumoniae* can undergo phase variation, switching to transparent variants that express lower capsule amounts and higher surface proteins (Weiser and Kapoor, 1999). Moreover, when in close contact with host cells, pneumococci reduce capsule levels, exposing adhesins that enhance interaction (Hammerschmidt, 2006). *S. pneumoniae* also employs exoglycosidases such as neuraminidase (NanA, NanB, and NanC), β-galactosidase (BgaA), and StrH, to modify host carbohydrate structures, thereby promoting adhesion (King, 2010). Recently, the concerted action of neuraminidases and pneumolysin in the destruction of the platelets was shown. Pneumococcal neuraminidases desialylate platelet glycoproteins, thereby increasing binding of pneumolysin, which finally leads to high cytotoxicity even at low toxin concentrations (Fritsch et al., 2024). Furthermore, pneumococci exploit host proteases such as plasmin, to degrade mucosal and extracellular matrix (ECM) components, facilitating stronger interactions with host cells (Bergmann and Hammerschmidt, 2007). Key surface adhesins including PspC, PavA, PavB, enolase, and pili allow *S. pneumoniae* to bind to host receptors and ECM proteins, promoting attachment and colonisation (Weiser et al., 2018; Jensch et al., 2010; Elm et al., 2004). PavA and PavB target and bind to different regions of fibronectin to enhance adhesion to host cells (Kanwal et al., 2017).

Lipoproteins such as PnrA, DacB, and MetQ, along with surface proteins like PspA, PspC, and PsaA, are promising candidates for vaccine development due to their role in adhesion and immune evasion (Narciso et al., 2024; Paulikat et al., 2024; Voß et al., 2018; Kohler et al., 2016). Additionally, the Pht proteins, especially PhtD and PhtE, are crucial for adherence and colonisation, making them valuable targets for therapeutic intervention (Khan and Pichichero, 2012; Kallio et al., 2014).In addition to the proteins mentioned earlier, proteases and peptidases also play a crucial role in bacterial adaptation to host environments and immune evasion (Marquart, 2021). These enzymes enable *S. pneumoniae* to degrade host tissues, modulate immune responses, and survive in nutrient-limited conditions commonly encountered during infection (Ali et al., 2021b; Bergmann and Hammerschmidt, 2006). Proteomic analyses have identified key enzymes whose abundance fluctuates during bacterial growth, potentially influencing the bacterium’s ability to transition between colonisation and invasive phases (Lee et al., 2006). The regulation of enzyme abundances under different growth conditions, particularly during stress, highlights the significance of these enzymes in the pathophysiology of *S. pneumoniae* (Lee et al., 2006).

Due to the limitations of infection-related and nutrient-defined *in vitro* media suitable to study the pathophysiology of pneumococci, we developed an optimized and user-friendly minimal medium in the present study. We then utilised this medium to elucidate the global proteome landscape of the non-invasive strain *S. pneumoniae* EF3030.

## 2 Methodology

### Bacterial strains and growth conditions

The bacterial strains used in this study (Supplementary Table S1) were initially retrieved from glycerol stocks and cultured on blood agar plates (BA), consisting of Columbia agar supplemented with 5% sheep blood. Following this, the strains were re-streaked onto fresh BA plates and incubated for a maximum of 8-10 hours at 37°C in the presence of 5% CO₂. Bacteria from these BA plates were then used to inoculate the liquid medium.

To compare growth, bacterial strains were grown in RPMI-based chemically-defined medium (CDM) as established by (Schulz et al., 2014) to ensure reproducible growth, addressing its limitations in supporting proper growth for certain *S. pneumoniae* serotypes. The medium was further modified by supplementing it with methionine, iron and manganese, resulting in CDM+ (Table 1 and 2).

**TABLE 1:** Stock solutions for modified CDM medium.

| Solutions | Components | Concentration [g/l] | Remarks |
| --- | --- | --- | --- |
| Buffer solution | NaHCO<sub>3</sub> | 24.7 | sterile filter and store at 4°C |
|  | Glycine | 1.11 |  |
|  | Choline chloride | 0.456 |  |
|  | NaH<sub>2</sub>PO<sub>4</sub> · H<sub>2</sub>O | 3.195 |  |
|  | Na<sub>2</sub>HPO<sub>4</sub> · 2H<sub>2</sub>O | 9.22 |  |
|  | Glucose | 74 |  |
| Adenine/Uracil solution | Adenine | 4 | dissolve Adenine and Uracil in 40ml 1M HCl at 90°C, then adjust the total volume with distilled water, sterile filter and store at room temp. |
|  | Uracil | 8 |  |
| Iron and Manganese solution | MnSO<sub>4</sub> · H<sub>2</sub>O | 0.7 | dissolve in sterile distilled water and store at 4°C |
|  | FeSO<sub>4</sub> · 7H<sub>2</sub>O | 0.625 |  |
|  | Fe(NO<sub>3</sub>)<sub>3</sub> · 9H<sub>2</sub>O | 0.125 |  |

**TABLE 2:**
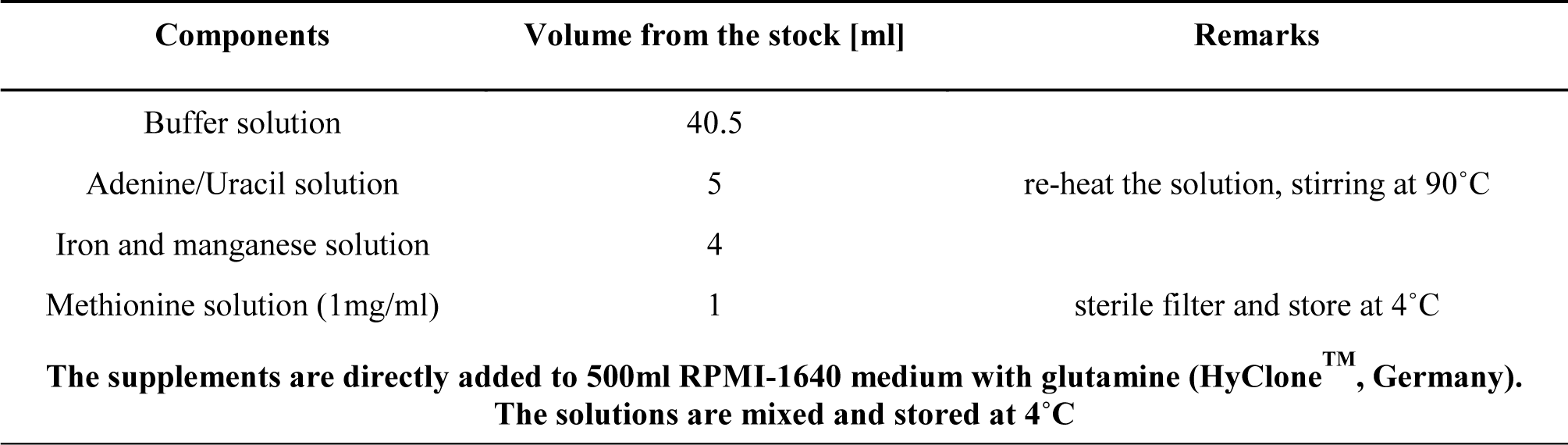
Preparation of modified RPMI 1640-based CDM+ medium.

For *S. pneumoniae, S.suis, S. mutans* and *S. agalactiae* pre-cultures were inoculated with an initial optical density (OD_600nm_) of 0.2 in CDM or CDM+. The strains were then sub-cultivated in fresh, pre-warmed minimal medium when they reached an optical density of 0.4, ensuring highly reproducible growth conditions during experiments. *S. pyogenes* was cultivated without subculturing, as the culture did not reach an optical density (OD) of 0.4. The growth of the other bacterial strains was monitored from an initial OD_600nm_ 0.2. For electron microscopic analysis, cultivation of pneumococcal strains was done in Todd-Hewitt broth with 0.5% yeast extract (THY). The strains were inoculated at an optical density (OD_600nm_) of 0.08.

### Field emission scanning electron microscopy (SEM) of pneumococcal strains

Pneumococcal strains were cultured in THY, CDM or CDM+ as described above, and samples were collected at an OD_600nm_ of 0.4 to 0.6. Bacteria were harvested by centrifugation at 3500 rpm, washed with phosphate-buffered saline (PBS), and fixed with 2.5% glutaraldehyde and 2.0% paraformaldehyde in washing buffer (0.1 M cacodylate buffer [pH 7], 0.09 M sucrose, 0.01 M CaCl_2_, 0.01 M MgCl_2_) for 1 hour at 4°C. Following fixation, pneumococci were immobilized to poly-L-lysine-coated coverslips for 90 minutes at room temperature, and fixation continued at 4°C overnight. The samples were then washed three times with washing buffer for 5 minutes each time, followed by treatment with 1% osmium tetroxide in washing buffer for 1 hour at room temperature. After three additional washing steps with washing buffer for 5 minutes each time, the samples were dehydrated in a graded series of aqueous ethanol solutions (10%, 30%, 50%, 70%, 90%, 100%) on ice for 15 minutes at each concentration. Prior to the final change to 100% ethanol, the samples were allowed to reach room temperature and were then critical point-dried using liquid CO_2_. Finally, the samples were mounted onto aluminium stubs, sputter-coated with gold/palladium, and examined with a field emission scanning electron microscope Supra 40VP (Carl Zeiss Microscopy Deutschland GmbH, Oberkochen, Germany) using the Everhart-Thornley SE detector and the inlens detector at a 80:20 ratio at an acceleration voltage of 5 kV. All micrographs were edited using Adobe Photoshop CS6 (Figure 2, Supplementary Figure S1).

### Preparation of *S. pneumoniae* cellular and supernatant proteins for proteome studies

#### Sample harvesting

*S. pneumoniae* strain EF3030 (serotype 19F) was cultured in CDM+ with sub-cultivation as described above. Pneumococci were harvested at three defined optical densities: early exponential phase (OD_600_ 0.4), mid exponential phase (OD_600_ 0.6), and late exponential phase (OD_600_ 1.0). A total of 6 mL of culture was centrifuged at 3900 rpm for 10 minutes, and the supernatant was carefully collected and the tube was immediately frozen by immersion in ice-cold ethanol and maintained at -70°C. The bacterial pellet was washed once with PBS, and the optical density was adjusted to 1.0. Subsequently, 1 mL of the bacterial suspension was centrifuged again at 3900 rpm for 5 minutes, and the resulting bacterial pellet was stored at -70°C until sample preparation.

#### Sample preparation

The collected samples of bacterial pellets were each resuspended in 50 µL of 20 mM HEPES buffer (pH 8) containing 1% SDS. Cell disruption was achieved through mechanical disruption using a bead mill at 3000 rpm for 3 minutes in liquid nitrogen. The resulting frozen powder was then dissolved in 150 µL of 20 mM HEPES buffer (pH 8) with 1% SDS. The solution was subjected to shaking at 1400 rpm at 95°C for 1 minute. After cooling, Pierce™ Universal Nuclease (Pierce, Thermo Fisher Scientific, MA, United States; 2.5 U, 4 mM MgCl2) was added, followed by an incubation in an ultrasonic bath for 5 minutes. Thereafter, all samples were centrifuged at 17000 g for 30 minutes, and the supernatant was carefully collected for subsequent analysis by mass spectrometry.

Proteins from the supernatant were precipitated using trichloroacetic acid (TCA). A volume of 1.5 mL of the supernatant was mixed with 225 µL of 100% TCA, resulting in a final concentration of 15% TCA. The solution was thoroughly mixed and incubated at 4°C for 48 hours. Following incubation, the solution was centrifuged at 17,000 g for 60 minutes at 4°C. The supernatant was carefully discarded, and the pellet was washed with 70% ice-cold ethanol by centrifugation at 17,000 g for 10 minutes at 4°C. This wash procedure was repeated three times with decreasing volumes of ethanol: 500 µL wash 1), 200 µL (wash 2), 180 µL (wash 3). After the final washing step, the ethanol was removed and the pellet containing the proteins was air-dried. The proteins were then resuspended in 15 µL of 20 mM HEPES buffer (pH 8) containing 1% SDS to solubilize the membrane proteins and incubated at 65°C for 2 minutes.

Protein concentrations of protein samples were determined using the Thermo Micro BCA Kit employing the BCA assay analysis pipeline included in the MassSpecPreppy pipeline (Reder et al., 2024).

Tryptic digestion of the pellet and supernatant-derived proteins was performed using the SP3 protocol (Reder et al., 2024). The resulting peptides were analysed by mass spectrometry on an Orbitrap Exploris^TM^ 480 mass spectrometer (Thermo Fisher Scientific), coupled to an Ultimate 3000 nano-LC system (Thermo Fisher Scientific). A data-independent acquisition (DIA) mode was utilised to acquire the data, for specific details see Supplement Table S3 and S4. Data are available via ProteomeXchange with identifier PXD062379 (Deutsch et al., 2023).

Reviewer Access via the PRIDE website (https://www.ebi.ac.uk/pride/) using the following details (Perez-Riverol et al., 2024; Perez-Riverol et al., 2016):

*Project accession: PXD062379*

*Token: aIkOo8bG7wCK*

#### Data analysis

The mass spectrometric data were analysed using the SpectronautTM Software (v19.1), via a spectral library-based approach. The search against a protein sequences database of *S. pneumoniae* EF3030 (<u>NZ_CP035897</u>) included tryptic peptides with up to two missed cleavages, fixed carbamidomethylation of cysteine and oxidation of methionine and N-terminal acetylation as variable modifications. Further analysis was performed using R version 4.4.1 and the SpectroPipeR package 0.3.0 (Michalik et al., 2025). The PneumoWiki (https://pneumowiki.med.uni-greifswald.de) annotations and descriptions of *S. pneumoniae* EF3030 proteins were used to analyse the genome’s functional response to the minimal medium.

For comparative analysis of protein composition between the cellular and supernatant fractions, intensity-based absolute quantification (iBAQ) values were calculated using Spectronaut. The relative iBAQ values were expressed as a percentage of the total sum of iBAQ values for each sample condition (Schwanhäusser et al., 2011) (Supplementary Figure S3). Protein localization were predicted using DeepLocPro v1.0 (Moreno et al., 2024) with ‘--group positive’ (Figure 4). Gene set enrichment analyses (GSEA; Subramanian et al., 2005) was performed using the fast gene set enrichment analysis (fgsea) R-package (Korotkevich et al., 2016), leveraging TIGRFAM (Haft et al., 2013) functional annotations of *S. pneumoniae* EF3030 proteins as retrieved from PneumoWiki and integrating orthologously mapped RegPrecise (Novichkov et al., 2013) based regulon information of *S. pneumoniae* TIGR4 (Supplementary Figure S2). Hierarchical clustering of protein profiles of proteins identified with at least two peptides in both fractions was performed in R using the hclust function (Figure 5, Supplementary Table S7). As distance metric, spearman correlation was applied and 1-r was used. For distance determination between clusters, the average method was used. Clusters cutting height was set to 40% of the maximal distance. Functional characterization of the clusters was performed by over representation analysis using the Fisher’s Exact test and employing TIGRFAM functional annotation and regulon information.

## 3 Results

### Methionine and the trace elements iron and manganese enhanced streptococcal fitness and growth in minimal medium

Previous studies have shown that amino acids are essential for improved growth behaviour in minimal medium for the cultivation of *S. pneumoniae,* serotype 2, D39 (Härtel et al., 2012). Later a RPMI based chemical-defined minimal medium was established with supplementation of uracil-adenine solution (CDM) (Schulz et al., 2014). However, *S. pneumoniae* EF3030, serotype 19F, and TIGR4, serotype 4, exhibited a prolonged lag phase and a lower yield in this medium (Ali et al., 2021a). To further improve the growth conditions and ensure reproducibility of pneumococcal growth, we supplemented CDM with methionine as well as with the metal ions, iron and manganese, which are needed as trace elements for bacterial growth. This newly developed minimal medium (CDM+) significantly reduced the lag phase of the strain EF3030. Similarly, growth of the other laboratory strains TIGR4, D39 and capsule knockout strains, D39Δ*cps* and TIGR4Δ*cps* was enhanced in CDM+ (Figure 1A and 1B). Furthermore, we cultivated pneumococcal clinical isolates of serotype 8, 12F, or 22F in CDM+ and monitored their growth. Indeed, a shorter lag phase and accelerated growth were observed for all strains (Figure 1C and 1D). To test whether this medium is also suitable for cultivating other streptococcal species, we cultivated *S. mutans, S. suis*, *S. pyogenes* (GAS) and *S. agalactiae* (GBS) in CDM+ (Figure 1E and 1F). However, *S. mutans* and *S. suis* strains showed similar growth in both media, CDM and CDM+ (Figure 1E). Remarkably, *S. pyogenes* did not grow at all in CDM and only marginally in CDM+, suggesting that this species may require additional components for growth in a minimal medium. In contrast, *S. agalactiae* grew to higher optical density in CDM+ (Figure 1F). Based on these results, we conclude that the improved minimal medium CDM+ can be applied to study the (patho-)physiology of different streptococcal species under reliable growth conditions.

**Figure 1:**
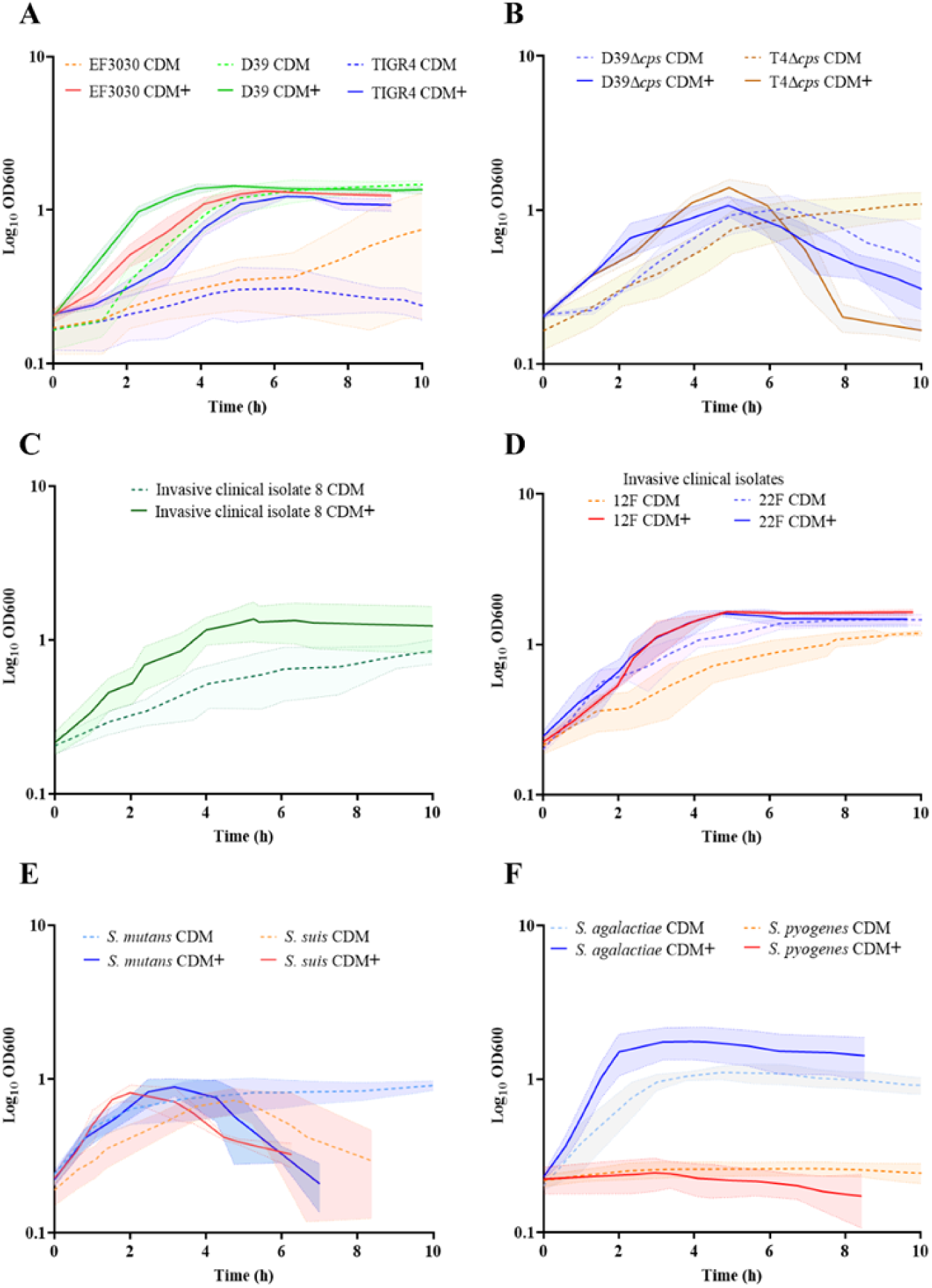
Optimization of a minimal medium for growth of pneumococcal and other streptococcal species. To establish a reliable and reproducible growth medium for pneumococcal and other streptococcal species, we compared the growth of these bacteria in two different media. Bacteria were cultured in RPMI supplemented with adenine and uracil (CDM), represented by the dotted lines, and in RPMI supplemented with adenine, uracil, methionine, iron, and manganese (CDM+), represented by the bold lines. The standard deviation (SD) of the measurements is represented by the shadow, indicating the variability of the results across three independent replicates (n=3). (A) Growth of pneumococcal strains, serotype 19F (EF3030), serotype 2 (D39) and serotype 4 (TIGR4) (B) Nonencapsulated mutants: D39Δ*cps* and TIGR4Δ*cps* (C) Invasive clinical isolates of *S. pneumoniae*: serotype 8 (D) Invasive clinical isolates of *S. pneumoniae*: 12F and 22F (E) *S. mutants* and *S. suis* (F) Group A streptococcal strain, *S. pyogenes* and Group B strain, *S. agalactiae*.

### Morphological characteristics of pneumococci in the exponential growth phase

Three encapsulated pneumococcal strains (D39, TIGR4, and EF3030) of different serotypes (2, 4, and 19F respectively) along with the isogenic non-encapsulated mutants (TIGR4Δ*cps* and D39Δ*cps*) were cultivated in three different media: the complex medium THY, as well as the minimal media CDM and CDM+. Pneumococci were harvested in exponential growth phase at an optical density of (OD_600nm_) 0.6 and prepared for scanning electron microscopy to visualize and illustrate the organization and morphology of the pneumococcal strains at high resolution. The encapsulated strains EF3030, TIGR4, and D39 formed aggregates in CDM+, whereas they did not form these assemblages when cultivated in either CDM or THY (Figure 2, Supplementary Figure S1), suggesting that the bacterial aggregates are due to the supplements present in CDM+. In contrast to wild-type pneumococci, the isogenic non-encapsulated strains TIGR4Δ*cps* and D39Δ*cps* appear smooth due to the lack of capsule and did not form substantial aggregates even in CDM+, suggesting an impact of the capsular polysaccharide (CPS) on pneumococcal aggregation. Notably, the encapsulated wild-type pneumococci were also surrounded by some vesicle-like structures that provided a rough texture to the bacterial surface, which was predominantly visible in CDM+ and appeared to contribute to aggregation (Figure 2, Supplementary Figure S1). Furthermore, serotype 4 strain TIGR4, which is an invasive clinical isolate, formed longer chains in the complex THY medium compared to the minimal media. TIGR4 wild-type strain also displayed pili-like structures on the surface and mostly near the septum region.

**Figure 2:**
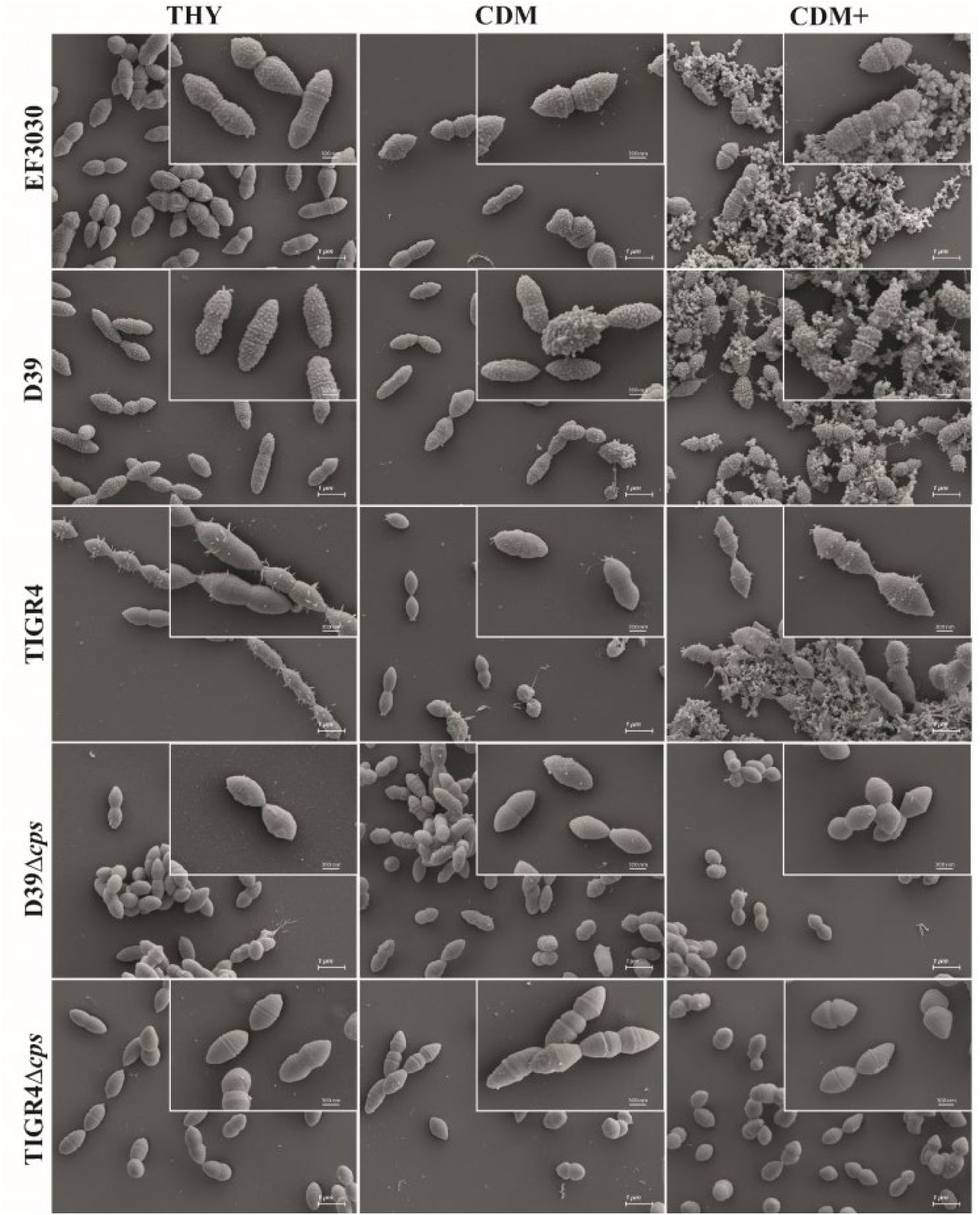
Scanning electron micrographs of *S. pneumoniae* cultivated in THY, CDM, or CDM+. EF3030 (serotype 19F), D39 (serotype 2), TIGR4 (serotype 4), D39Δ*cps* and TIGR4Δ*cps* were cultured at 37°C up to an optical density of OD_600nm_ 0.6 in each medium. Pneumococci were harvested and subjected to scanning electron microscopy. The micrographs were taken at a magnification of 10,000x and the inset micrographs in the top right corner were taken at a higher magnification of 30,000x. These higher magnifications provide more detail of the pneumococcal surface structures and aggregates. Scale bars = 1 µm; Scales bars of the inset micrographs = 300 nm (see Supplementary Figure S1).

### Global proteome analysis of *S. pneumoniae* EF3030, serotype 19F

To elucidate the (patho-)physiology of *S. pneumoniae* strain EF3030 at protein level when cultured in the minimal medium CDM+, we conducted a comprehensive proteomic analysis of both the cellular and the supernatant fractions. Although the genome of serotype 19F strain, EF3030 has been published (Junges et al., 2019), a detailed proteome profile is not yet available for this strain nor for any other serotype 19F strain. Our goal was to characterize the protein composition of *S. pneumoniae* strain EF3030 across different growth phases (early exponential, mid-exponential, and late exponential phases) using mass spectrometry.

#### Experimental Setup and Protein Identification

*S. pneumoniae* EF3030 was cultured in CDM+ and harvested at three growth phases, defined by optical density (OD600nm) values of 0.4 (early exp), 0.6 (mid-exp), and 1.0 (late exp) (Figure 3A). Total protein was isolated from both the cytosolic and supernatant fractions and proteome analysis was performed using the Pneumowiki annotated genome of strain EF3030 (NCBI RefSeq accession no. NZ_CP035897.1 (alternatively: CP035897). NZ_CP035897.1 predicts 1862 protein-coding genes, we were able to successfully identify 1381 proteins in the cytosolic fraction and 1373 proteins in the supernatant fraction, with more than two peptides with ion q-values less then 0.001 (Figure 3B). These numbers represent 74.16% and 73.73% of the annotated theoretical proteome in the cytosolic and supernatant fractions, respectively. The high number of different proteins in the supernatant indicated partial cell lysis already at an early stage. A comparison revealed 1,344 proteins common to both fractions, with 37 and 29 proteins uniquely identified in the cytosol and supernatant, respectively (Figure 3B).

**Figure 3:**
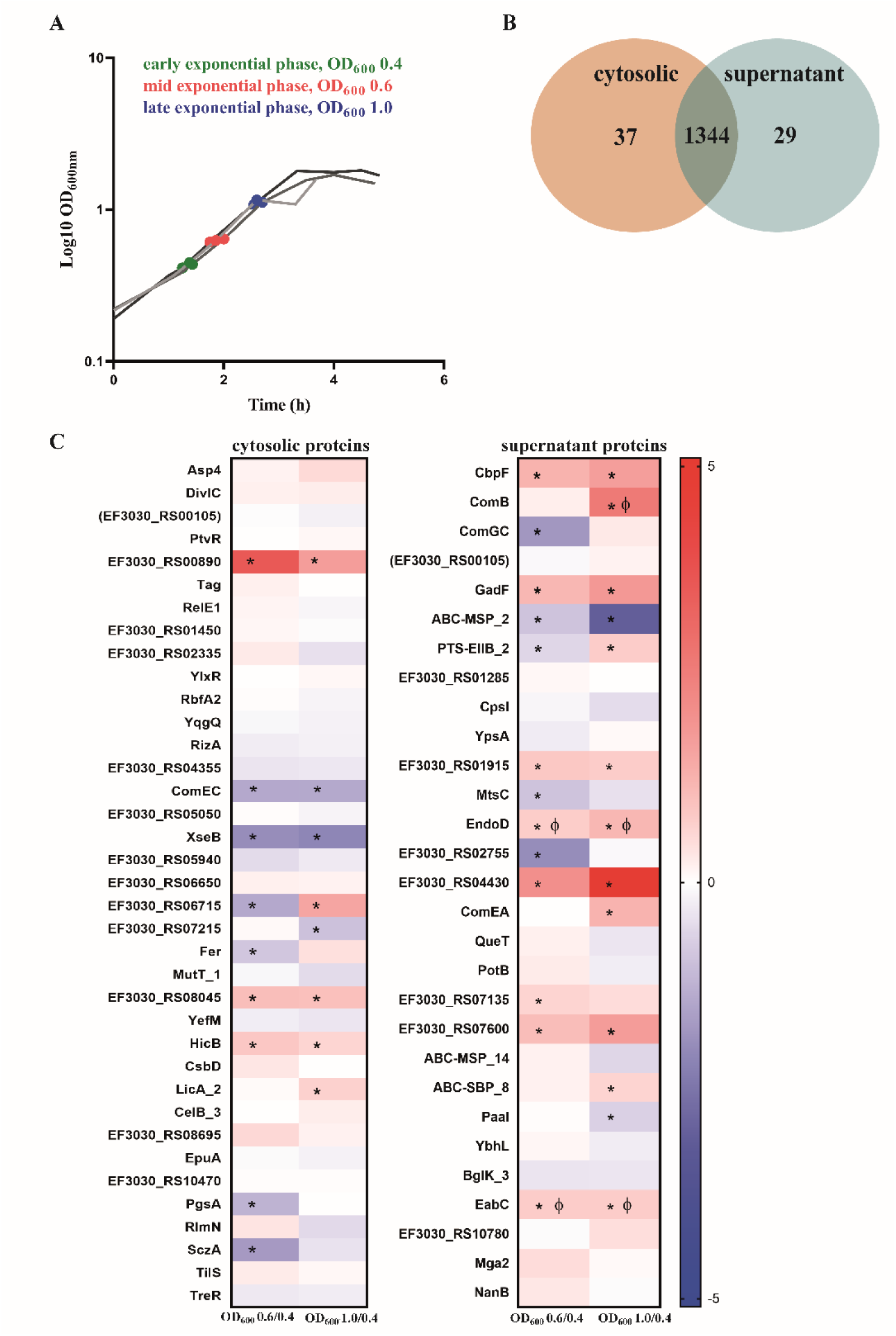
Identification of pneumococcal proteins exclusively detected in the cytosol or supernatant fractions. (A) Sampling of bacterial cultures and their corresponding supernatants were done at different exponential growth phases—early (OD₆₀₀ = 0.4, green), mid (OD₆₀₀ = 0.6, red), and late (OD₆₀₀ = 1.0, blue). The three lines represents the biological replicates. (B) A total of 1,381 proteins were identified in the cytoplasm and 1,373 proteins were identified in the supernatant, with two or more than two peptides analysed by mass spectrometry. Of these, 37 proteins were exclusively found in the cytosol, whereas 29 proteins were uniquely present in the supernatant, as depicted in the Venn diagram. (C) Abundance of the proteins detected exclusively in the cytosol and the supernatant. Heatmaps represent the signal log ratio comparisons during early, mid and late exponential growth phases. Φ indicates significant values, p-value less than 0.05. * Indicates absolute fold change greater than or equal to 1.5. The protein EF3030_RS00105 is in ‘brackets’ because the identified peptides could originate from other annotated proteins.

#### Proteins Identified in the Cytosolic Fraction

In the cytosolic fraction, 37 proteins were uniquely identified (Figure 3C), including key stress response and growth-regulatory proteins. ComEC, which is involved in DNA internalisation related to competence and PtvR, important for fitness coping with antibiotic and stress, were detected in the cytosol (Pimentel and Zhang, 2018; Liu et al., 2017). Notably, the toxin-antitoxin system components YefM, RelE1, and HicB, which are implicated in bacterial persistence, were identified in this fraction only. Bacterial persistence is a crucial factor in the survival of *S. pneumoniae* within the host and during antibiotic treatment. HicB was of particular interest, as it exhibited an 1.7 fold increase during the mid-exponential phase and 1.5 fold during the late exponential phase (Supplementary Table S5).

#### Proteins Identified in the Supernatant Fraction

The supernatant fraction contained 29 unique proteins (Figure 3B). Most of these proteins were truly secreted or surface associated, including CbpF, ComGC, ComB, ComEA, NanB, EndoD, MtsC, QueT, CpsI, and EabC (Figure 3C). However, we also detected some cytoplasmic proteins such as Mga2, a virulence factor transcriptional regulator, together with Paal, and YbhL, mainly due to a substantial lysis rate. Notably, a significant proportion of the detected proteins in the supernatant fraction contribute directly or indirectly to virulence, including NanB, CpsI, ComEA and EndoD. These proteins exhibited distinct abundance patterns during growth, with EndoD and ComEA increasing in later growth phases, whereas CpsI showed a decrease in abundance under the same conditions. Furthermore, EabC, associated with a domain that could be linked to host interaction and the competence factor ComB displayed significant increases in abundance during the late exponential phase. The choline-binding protein CbpF also displayed a notable 3.2 fold increase during the late log phase. The surface proteins, including CpsI and ComEA, demonstrated phase-dependent abundance fluctuations, reflecting the adaptive response of *S. pneumoniae* to changing growth conditions.

#### Protein Composition Comparison: Cytosol vs. Supernatant

The comparative analysis of the relative iBAQ values for cytosolic proteins and proteins present in the supernatant revealed a wide range of protein abundances, spanning approximately 6 log10 steps (Supplementary Figure S3A). Highly abundant proteins, including Gap (GAPDH), Tuf, enolase (Eno), EF3030_RS05545 (PtsH), EF3030_RS02365 (Pgk) and ribosomal proteins (e.g., RplX), were found in both fractions. Notably, when the total proteins detected in the late exponential phase were analysed, LytN and SpxB were found among the top 10 highest abundant proteins. LytN, a murein hydrolase, plays a crucial role in cell division and cell wall integrity. Overexpression of LytN has been shown to cause cell lysis in other bacteria like *S. aureus* (Frankel et al., 2011). SpxB, a pyruvate oxidase, is key enzyme for H₂O₂ production (Bättig and Mühlemann, 2008). The protein composition in the supernatant fraction in general mirrored that of the cytosolic profile (Supplementary Figure S3B). Proteins that were at least 100-fold enriched in the supernatant fraction, including those involved in cell surface attachment and extracellular functions, such as PfbA, ZmpD, ZmpB, MapP, EF3030_RS10100, EF3030_RS00555, and PcsB. Conversely, we identified the ribosomal protein RpmC, EF3030_RS06250 (AroH), EF3030_RS02670, EF3030_RS02470, and EF3030_RS10365 as being at least 20-fold enriched in the cytosolic fraction.

*In silico* predictions of protein localisation by DeepLocPro (Moreno et al., 2024) indicate that the supernatant fraction is substantially enriched in extracellular proteins and cell wall and surface proteins across all analysed growth states (Figure 4). Notably, the relative abundance of membrane and cytoplasmic proteins does not differ between the two fractions, a finding that is consistent with the high frequency of lysis of pneumococcal cells (Terrasse et al., 2015;Eldholm et al., 2009)

**Figure 4:**
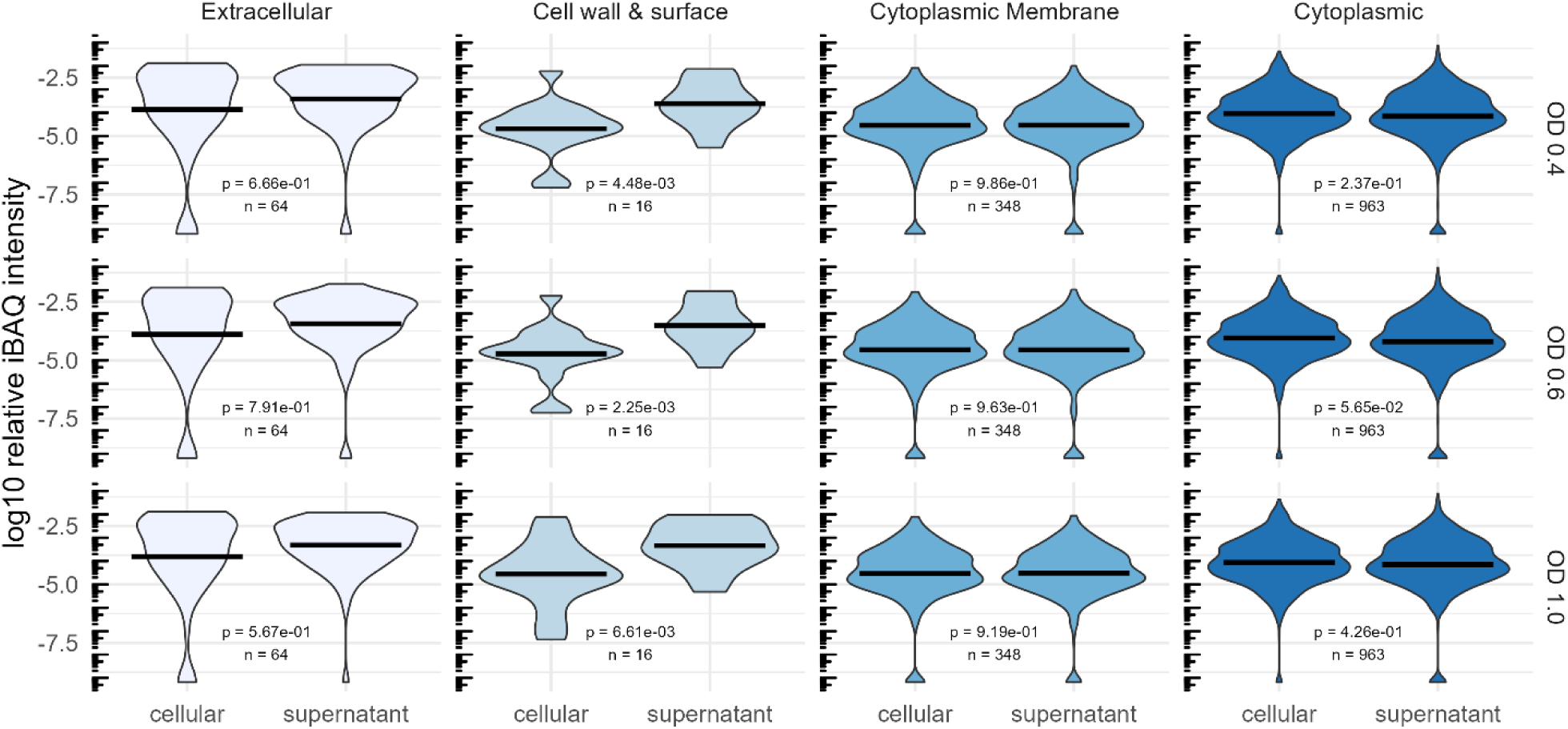
Comparison of the abundance of proteins regarding their predicted localization by DeepLocPro (Moreno et al., 2024). The violin plot shows log10-transformed relative iBAQ values. Medians are indicated as lines. Significant differences of mean relative abundances between the cellular and supernatant (secretome) fraction was tested using a Wilcoxon-test.

#### Growth-dependent changes of the cellular proteome and of the exoproteome

To investigate pneumococcal physiology in the newly developed minimal medium CDM+, pneumococcal cells and supernatants were collected at early, mid, and late exponential growth phases as shown in Figure 3A Comparative analysis of the mid and late exponential phase samples versus early exponential phase samples allowed for the examination of changes in protein expression profiles, metabolic alterations, and bacterial cell adaptations across varying growth states and cell densities (Figure 5). This information is crucial for the development of therapeutics targeting bacterial proteins, as it can provide insight into strategies to inhibit bacterial growth and progression of infection at different stages. The distinct growth phases correspond to different numbers of bacteria during colonization or dissemination in the blood or cerebrospinal fluid. Initially low bacterial number initiate adherence or cross host barriers and subsequent accumulation or biofilm formation contribute to the progression of infection (Hall-Stoodley et al., 2004).

**Figure 5:**
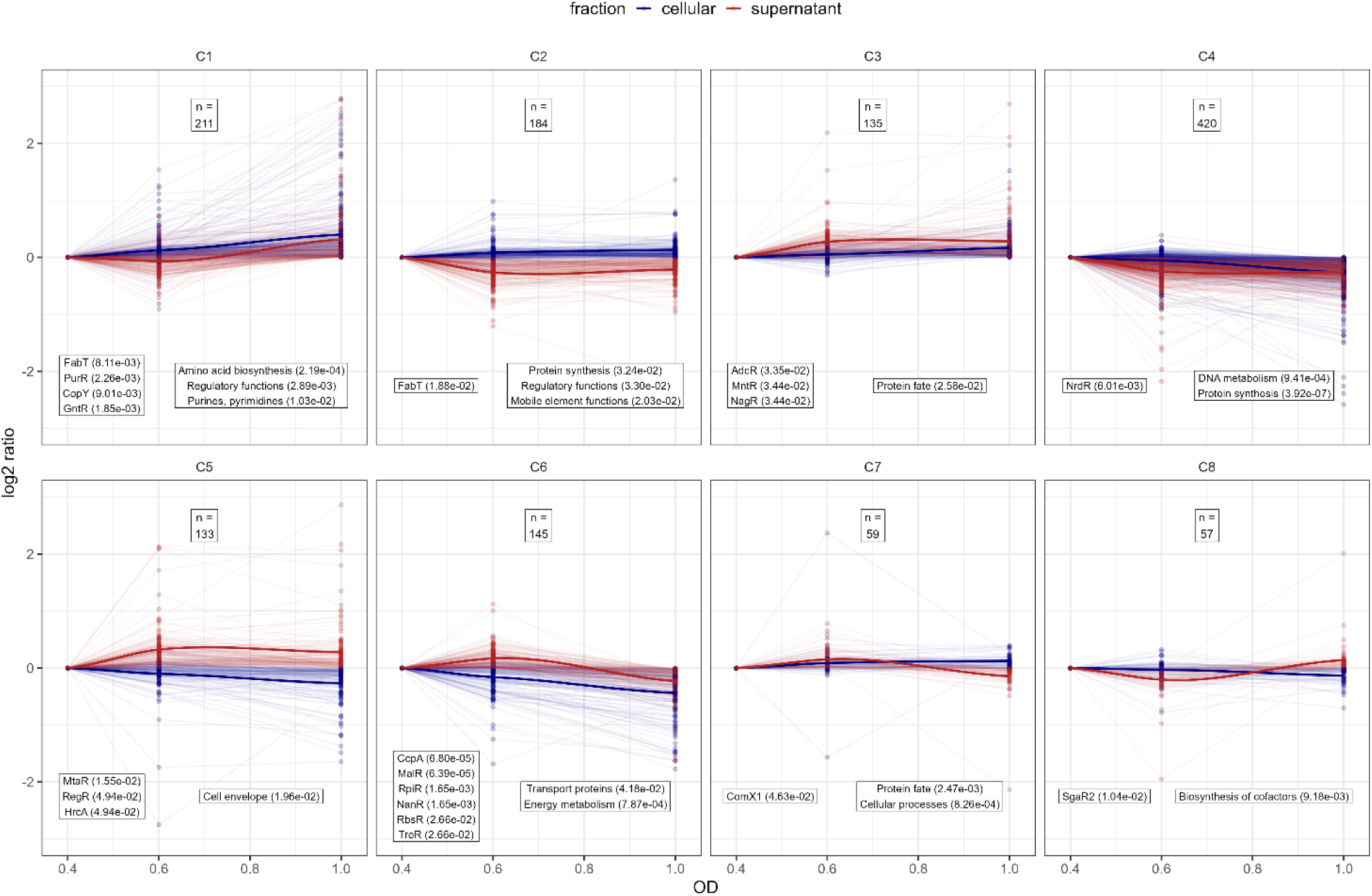
Analysis of protein profile clusters. For clustering the log2 protein intensity ratios compared to OD_600nm_ 0.4 were used for the supernatant and cellular fraction. Line plots of protein profiles for each cluster are shown, with cellular fraction profiles are depicted in blue and supernatant fraction profiles in red. To show the general cluster profile the LOESS-fit of all protein profiles are depicted as solid line (Jacoby, 2000). The total number of proteins in each cluster is indicated at the top of each cluster panel. Each cluster was tested for over-representation of proteins belonging to known regulons and known TIGRFAM functional annotations according to PneumoWiki (https://pneumowiki.med.uni-greifswald.de). Over-representation was assessed using one-sided Fisher’s exact test (p-value ≤ 0.05). Over-represented regulons and functional categories are depicted in the left box and right box, respectively, along with their corresponding p-values in brackets.

To gain a general overview about the protein abundance changes along the growth states, the obtained proteome profiles of *S. pneumoniae* EF3030 were normalized to the protein abundance at the early exponential phase and then clustered. Hierarchical clustering was applied (Supplemental Table S7 for the data) and clusters were defined by cutting of 40% of the total tree distance. This resulted in eight general protein clusters representing the proteome dynamics along the growth states in the supernatant and cytoplasmic fraction (Figure 5). Clusters C1 and C6 represent proteins associated with metabolic functions as identified using overrepresentation analysis of known regulons and functional TIGRFAM categories. Of note, proteins belonging to the fatty acid biosynthesis FabT regulon and the purine synthesis PurR regulon as well as amino acid biosynthesis proteins tend to accumulate along the growth state in the cytoplasmic and supernatant fraction, whereas proteins belonging to the central sugar metabolism CcpA regulon, the maltose utilization MalR regulon, sialic acid utilization NanR and RpiR regulons, fructooligosaccharides utilization RbsR regulon and the trehalose utilization TreR regulon decreased in abundance in the cytosolic and supernatant fraction during progression through growth phases (Figure 5). In that sense, our data reveal a shift from sugar-based glycolysis to other metabolic aspects over time when growth in the newly optimized medium and metabolic dynamics also shape the pathophysiological potential of pneumococci. In general, the CcpA regulated central metabolism is closely linked to virulence (Iyer et al., 2005; Carvalho et al., 2011; Zhang et al., 2023) and CodY-driven metabolic aspects are essential in colonization and infection processes of pneumococci (Hendriksen et al., 2007;Johnston et al., 2015). Proteins of the clusters C3 and C5 could be quantified in higher abundance in the later phases of growth in the supernatant. These clusters comprised of the zinc acquisition regulator AdcR, manganese homeostasis MntR regulon, the N-acetylglucosamine utilization and the hyaluronate utilization, RegR regulon as well as proteins function associated to the cell envelope, demonstrating the need of metal ions and accumulation of classic extracellular proteins during later growth phases. Proteins of the DNA metabolism, protein biosynthesis (cluster C4) and biosynthesis of cofactors (cluster C8) remained relatively stable during the exponential growth phase, highlighting their essential role in the bacterial physiology.

To gain detailed insights into the growth state dependent changes in protein levels in both the cytosolic and supernatant fraction, pairwise comparisons were performed for the samples harvested in the late exponential phase to samples harvested in early exponential phase using the ROPECA approach (Suomi and Elo, 2017) (Figure 6, Supplementary Table S5 and S6). Only proteins with an absolute fold change of 1.5-fold or greater and an adjusted p-value less than 0.05 were considered as significantly altered and were analysed (datasheet attached to the supplement, Table S4). 34 proteins showed increased abundance in both fractions, the cytosol and supernatant.

**Figure 6:**
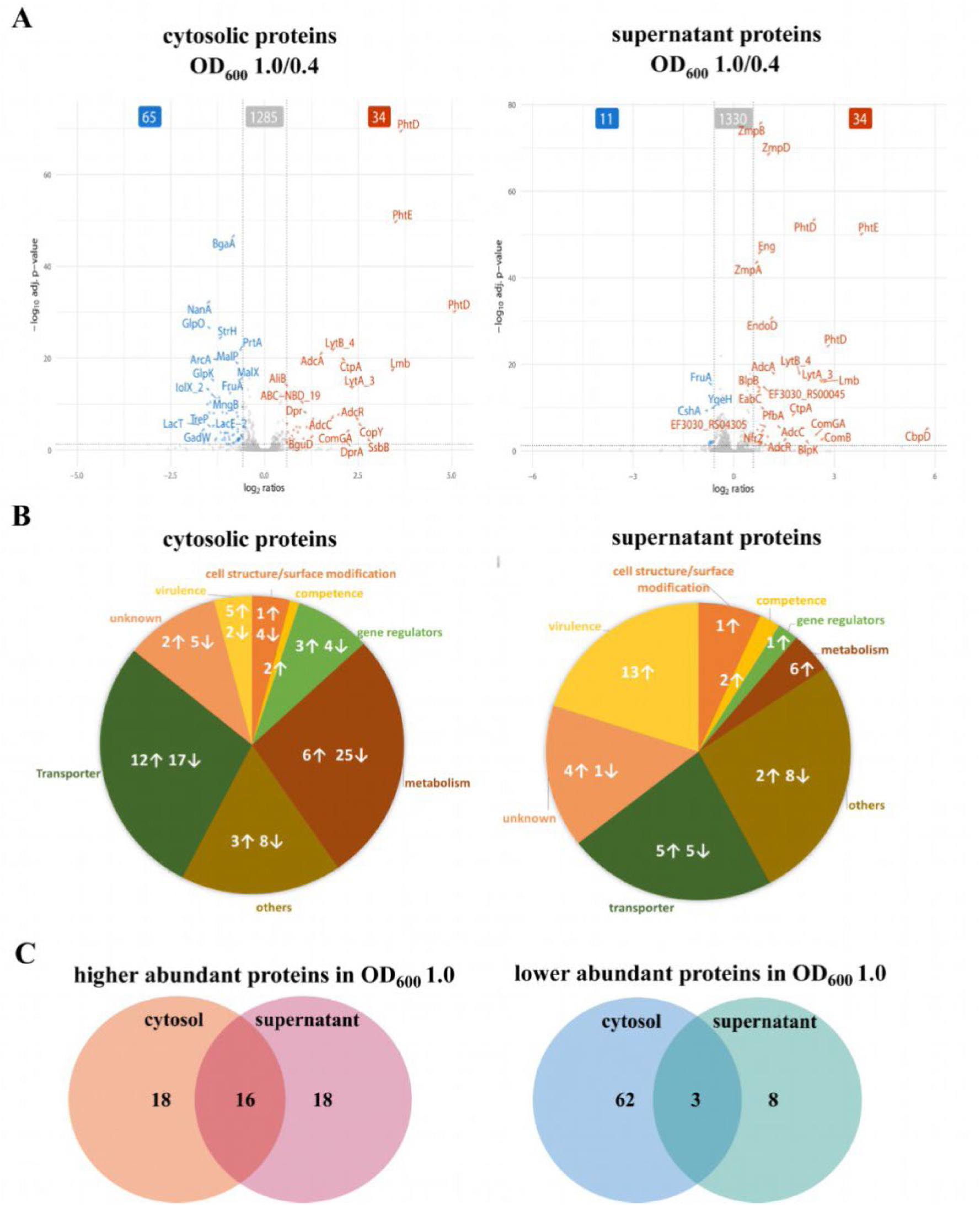
Quantification of the abundant proteins in the cytosol and the supernatant fractions during exponential growth. (A) Significant changes in protein abundance during the late exponential phase compared to the early exponential phase are visualized using volcano plots. The data represents the significant signal log ratio comparison between the different optical densities, with p-value less than 0.05 and a fold change greater than 1.5-fold. Proteins with higher abundance are labelled red, whereas proteins with a decrease in protein abundance are marked in blue. (B) Pie charts illustrate the significantly abundant proteins and their functions in the cytosol and supernatant during the exponential growth phase. Arrows indicate an increase or a decrease in protein abundance by comparing late log to early exponential phase. (C) A selection of proteins with increased abundance in the cytosol or supernatant, as well as proteins with reduced abundance in the cytosol or supernatant, at late growth.

In contrast, 65 proteins were significantly lower in abundance in the cytosol fraction and 11 proteins were significantly lower in the supernatant fraction (Figure 6A). We then classified these differentially expressed proteins into functional groups of interest: transporter proteins (29 in cytosol and 10 in supernatant), gene regulation proteins (7 in cytosol and 1 identified in supernatant), metabolic proteins (31 in cytosol and 6 in supernatant), virulence and competence factors (9 in cytosol and 15 in supernatant), proteins with others functions (11 in cytosol and 10 in supernatant) and proteins of unknown function (7 in cytosol and 5 in supernatant) (Figure 6B). Detailed description along with the fold change is shown in the Supplementary Table S4A and S4B.

To determine the overlap of the proteins shown in the volcano plot (Figure 6A) in both the cytosolic and the supernatant fraction, a Venn diagram (Figure 6C) was used. Sixteen proteins (the Pht family proteins, the Adc system proteins involved in Zinc homeostasis, ComGA, sortase SrtA, YdcP_2, BguA, Mip, Nfr1, Nfr2) were found to significantly increased in both the cytosol and the supernatant, while three proteins (FruA, TreP and GadW) were significantly reduced in both fractions (Supplement Table S4A and S4B, proteins marked with “*”).

#### Transporters

During the late-log phase (OD600 1.0), increased abundance of various transporter proteins was measured in both the cytosolic and supernatant fractions (Figure 6B, Supplementary Table S4A). Notably, PTS and ABC transporters involved in sugar uptake, such as BguB, BguC, and BguD, were more abundant in the cytosol. Moreover, zinc and metal transporters, such as AdcA, AdcAII, AdcC, and CtpA, were elevated in both fractions, indicating the bacterial adaptation to metal ion availability during growth, as indicated by overrepresentation of the AdcR regulon in cluster C3 (Figure 5). In contrast, several sugar transporters, including MalC, BrnQ, GadW, FruA, and TreP exhibited reduced abundance during the late-exponential phase (Supplementary Table S4B). This is in line with the observation that the general sugar metabolism decreases in later growth states (cluster C6; Figure 5).

#### Gene regulators

Several transcriptional regulators, including the Zn-dependent regulator AdcR showed increased abundance in both the cytosolic fraction and the supernatant fraction during the late-exponential phase (Figure 6A, Supplementary Table S4A). Other regulators like GntR and CopY were significantly increased in the cytosolic fraction during the later growth phase (Supplementary Table S5). In contrast, other regulators such as NmlR and FruR, exhibited a significant reduction in the cytosol and were also reduced in the supernatant during later growth phases suggesting growth-phase-specific regulation of bacterial adaptation and stress response (Supplementary Tables S4B, S5 and S6).

#### Metabolic proteins

Metabolic proteins involved in amino acid and sugar metabolism, such as galactokinase (GalK), LacG, and arginine deaminase system (ArcA, ArgF(ArcB), ArcC,), exhibited decreased abundance during the late-log phase in the cytosol (Supplementary Figure S2, Table S4B). In contrast, some metabolic enzymes, including GuaA2, PEP phosphomutase, YdpC_2, and BguA, showed increased abundance in the cytosol, indicating changes in metabolic activity during different growth phases (Supplementary Table S4A). This metabolic adaptation is also reflected by the identification of the two general metabolic clusters C1 and C6 (Figure 5). In general, sugar metabolism proteins tend to decrease during the late exponential phase in both the cytosolic and supernatant fractions, whereas the abundance of proteins involved in amino acid and purine and pyrimidine biosynthesis increased.

This shift in metabolic regulation is further supported by gene set enrichment analysis (GSEA), as depicted in Supplementary Figure S2. The plot illustrates functional annotation and regulon-based enrichment across different growth phases (OD 0.6/0.4 and OD 1.0/0.4). The colour of the dots represents the direction of normalized enrichment scores (NES), with red indicating positive enrichment and blue indicating negative enrichment. The size of each dot corresponds to the absolute NES value, highlighting the magnitude of enrichment. Notably, during the transition from mid to late exponential phase, cellular processes and nucleotide metabolism were positively enriched, while energy metabolism showed a negative enrichment. Additionally, the CcpA regulon, known for its role in carbon metabolism, exhibited significant negative enrichment, reinforcing the observed decline in sugar metabolism proteins (Supplementary Figure S2). These findings collectively suggest a metabolic shift favouring nucleotide and amino acid biosynthesis over sugar metabolism during later growth stages.

#### Competence and Other proteins

Remarkably, we observed a more than 2-fold increase of the competence type IV pilus ATPase ComGA in both pneumococcal fractions during the late exponential phase (Figure 6A, Supplementary Table S4A). In addition, ComB, a component of the competence system, was found to be 6.1 fold increased in the supernatant at the late exponential phase (Figure 6A, Supplementary Table S4A). Competence requires a complex and well-timed interplay of several regulons (such as ComE, ComX, BlpR) (Winkler and Da Morrison, 2019; Weyder et al., 2018) We found that the ComX1 regulon has a general peak in the mid exponential phase while protein levels decrease in the supernatant fraction during the late exponential phase (cluster C7; Figure 5)

Additionally, several cytoplasmic proteins involved in DNA processing and repair showed increased abundance over time (Figure 6A, Supplementary Table S4A). The single-strand DNA-binding protein, SsbB (EF3030_RS09295) stabilizes single-stranded DNA during replication, recombination, and repair. DprA facilitates natural transformation by binding and guiding incoming DNA into the chromosome. Sortase A, although primarily responsible for anchoring surface proteins, may support competence by attaching transformation-related proteins to the cell surface. SsbB and DprA are directly involved in DNA uptake and processing during natural transformation, a process that enhances genetic adaptability. Sortase A might contribute to this process by supporting the display of competence-associated factors on the cell surface. In the supernatant fraction, bacteriocin (EF3030_RS00555) or bacteriocin-like protein BlpU, bacteriocin secretion accessory protein (EF3030_RS02515), serine hydrolase (EF3030_RS00045), a glycoside hydrolase (EF3030_RS10670), and a surface protein (EF3030_RS05660) increased significantly at late exponential phase (Supplementary Table S4A). The secreted proteins (bacteriocins, hydrolases, and surface proteins) indicate enhanced bacterial competition, environmental adaptation, and potential host interactions. In contrast,out of the 11 significantly reduced proteins in the supernatant (Figure 6C), 8 proteins were exclusively found in the supernatant including YqeH, RsgA, ParB partition protein, methyltransferase domain containing protein, and the DEAD box helicase, whereas 3 were also found in the cytosol (Figure 6C, Supplementary Table S4B).

#### Protein profiles of surface proteins and proteases - investigation of the repertoire of selected virulence factors

For comparative analysis, we classified various proteins based on their surface anchoring mechanisms (Figure 7) (Lane et al., 2022; Bergmann and Hammerschmidt, 2006; Pérez-Dorado et al., 2012). The data revealed a significant increase in the signal log ratio of several lipoproteins, choline-binding proteins, sortase-anchored proteins, and non-classical surface proteins (NCSP) in the supernatant fractions comparing the mid and late log phase to the early log phase (Figure 7, Supplementary Table S5 and S6). Choline-binding proteins were detected in both the cytoplasmic and supernatant fractions, with the most striking increase observed for the murein hydrolase CbpD, which exhibited a 54.7-fold increase in abundance in the supernatant during the late log phase (Figure 7, Supplementary Table S4A). A 3.2-fold increase in abundance was also detected for CbpF, which was exclusively found in the supernatant (Figure 3). In contrast, while all choline-binding proteins were present in the cytoplasmic fraction at low levels, their abundance remained relatively unchanged throughout growth. We observed PhtD to be the most significantly elevated protein detected in both the cytoplasmic and supernatant fractions during the later stages of log growth (Figure 6A). PhtD and PhtE are members of the histidine triad protein (Pht) family, characterized by four to six histidine motifs that bind divalent metal cations, such as Zn²⁺ (Bersch et al., 2013).

**Figure 7:**
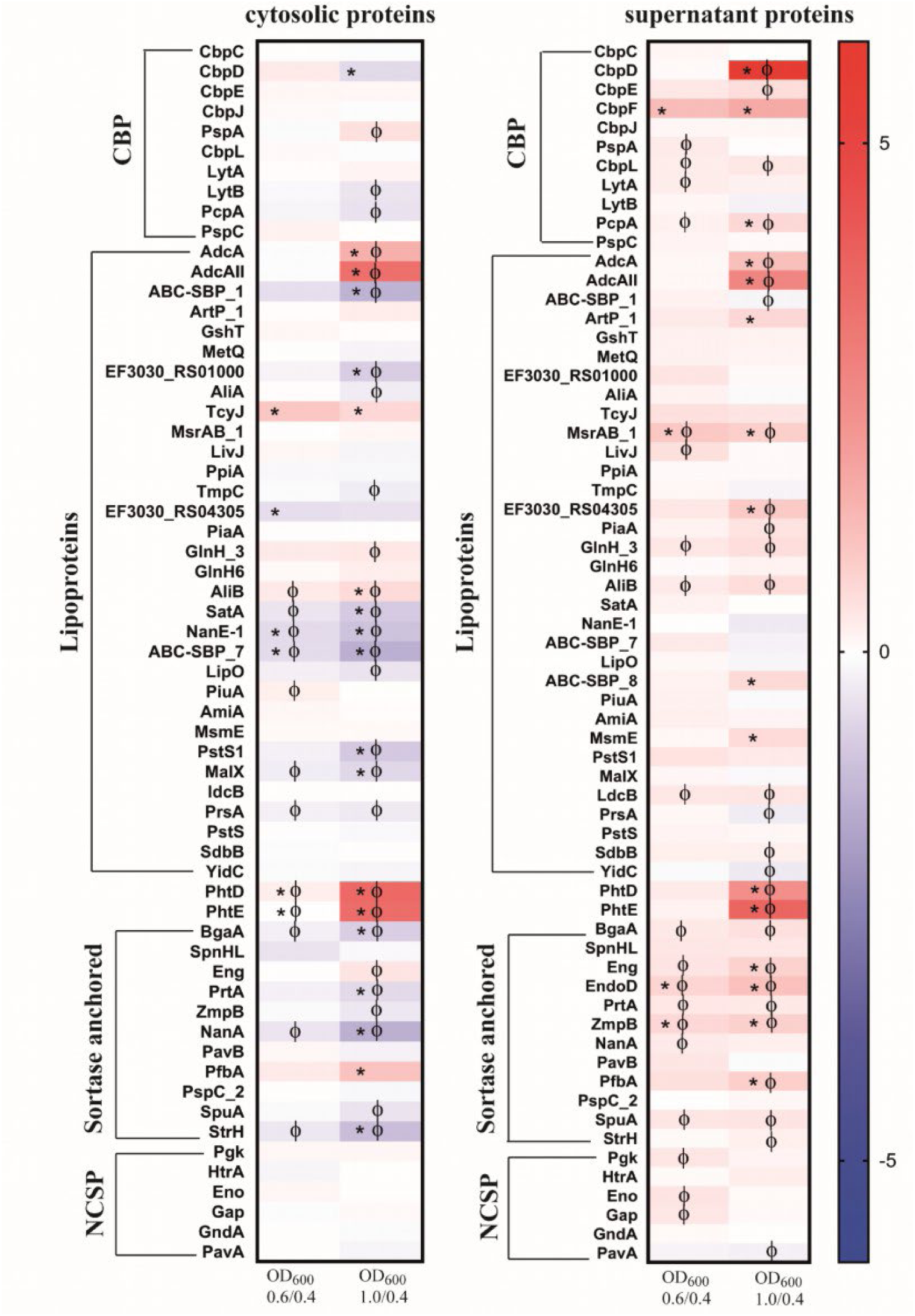
Heat maps of surface associated proteins with two or more peptides identified both in the cytosol and the supernatant of strain EF3030 by mass spectrometry analysis. The abundance of detected proteins was classified as choline-binding proteins (CBP), lipoproteins, sortase-anchored proteins and non-classical surface proteins (NCSP) in the cytosol and supernatant. The heat map represents the signal log ratio values between mid-exponential phase to early exponential phase and late exponential phase to early exponential phase. Φ indicates significant values, p-value less than 0.05. * Indicates an absolute fold change greater than or equal to 1.5.

Among the lipoproteins, AdcA and AdcAII were the prominent proteins with higher abundance, found in the late log phase in both the cytosolic fraction and extracellular fraction. A significant increase of ABC-SbP_8 was observed at the late exponential phase in the extracellular fraction, whereas other ABC-SBP (-SBP_1 and –SBP_7) components remained unchanged throughout growth. Furthermore, MsmE, ArtP_1, RS04305, and AliB as subunit of the Ami oligopeptide ABC transporter showed higher abundance in the late log phase. AliA, another subunit of the Ami transporter system, did not exhibit significant changes in abundance during growth. Notably, MsrAB, as part of the detoxification system for oxidized proteins, was consistently detected in higher abundance across all growth conditions compared to the early exponential phase. Additionally, several other lipoproteins like LdcB or SdbB remained stable during growth (Figure 7).

Significant differences in the abundance of sortase-anchored proteins compared to the early exponential phase were observed in the supernatant fraction. A notable increase in the abundance of EndoD, which contains an Ig-like domain, was found in the extracellular fraction. Additionally, increases were also observed for Eng, ZmpB, and PfbA. In contrast, the neuraminidase NanA level remained unchanged during growth, and similarly, SpnHL SpuA, StrH and PspC_2 did not exhibit alterations. A decreased abundance was detected for the adhesion PavB in the late exponential growth phase. Furthermore, decreases in NanA and StrH levels were observed in the cytoplasmic faction during exponential growth phases (Figure 7).

The non-classical surface proteins (NCSP) like enolase (Eno) and GAPDH (Gap) were found in abundance in both the cytosol and the supernatant (supplementary Table S4A). They were observed in the early exponential phase and significantly increased during the mid-exponential phase (Figure 7). Notably, PavA (pneumococcal adherence and virulence factor) lacking a secretion signal and domains for cell surface anchoring increased significantly during the late exponential phase in the supernatant fraction.

Proteases and peptidases are essential for multiple physiological processes in bacteria. Most of the proteases and peptidases discussed in this study (Figure 8) maintains important cellular functions attributing to survival, adaptability and pathogenesis (Marquart, 2021). The typical Clp protease components (ClpP, ClpX, ClpC and ClpE) and the chaperone ClpL, were found in both fractions. The ClpP ratio decreased in the cytoplasmatic fraction during the late exponential phase compared to the early growth phase, however there is no significant change in either fractions during the exponential growth. Interestingly, out of the three extracellular serine proteases present in the colonising strain, EF3030, the secreted serine proteases PrtA and HtrA were identified in both the fractions. No significant change of HtrA was observed during the exponential growth. In the cytosol, PrtA significantly reduced more than 1.5 fold during the late exponential phase, however it increased significantly in the supernatant during the exponential growth (Supplementary Table S4A). CbpG, the third serine protease was not detected under our experimental conditions. YdcP_2, a member of the U32 family peptidases, showed an increased abundance during growth in both fractions.

**Figure 8:**
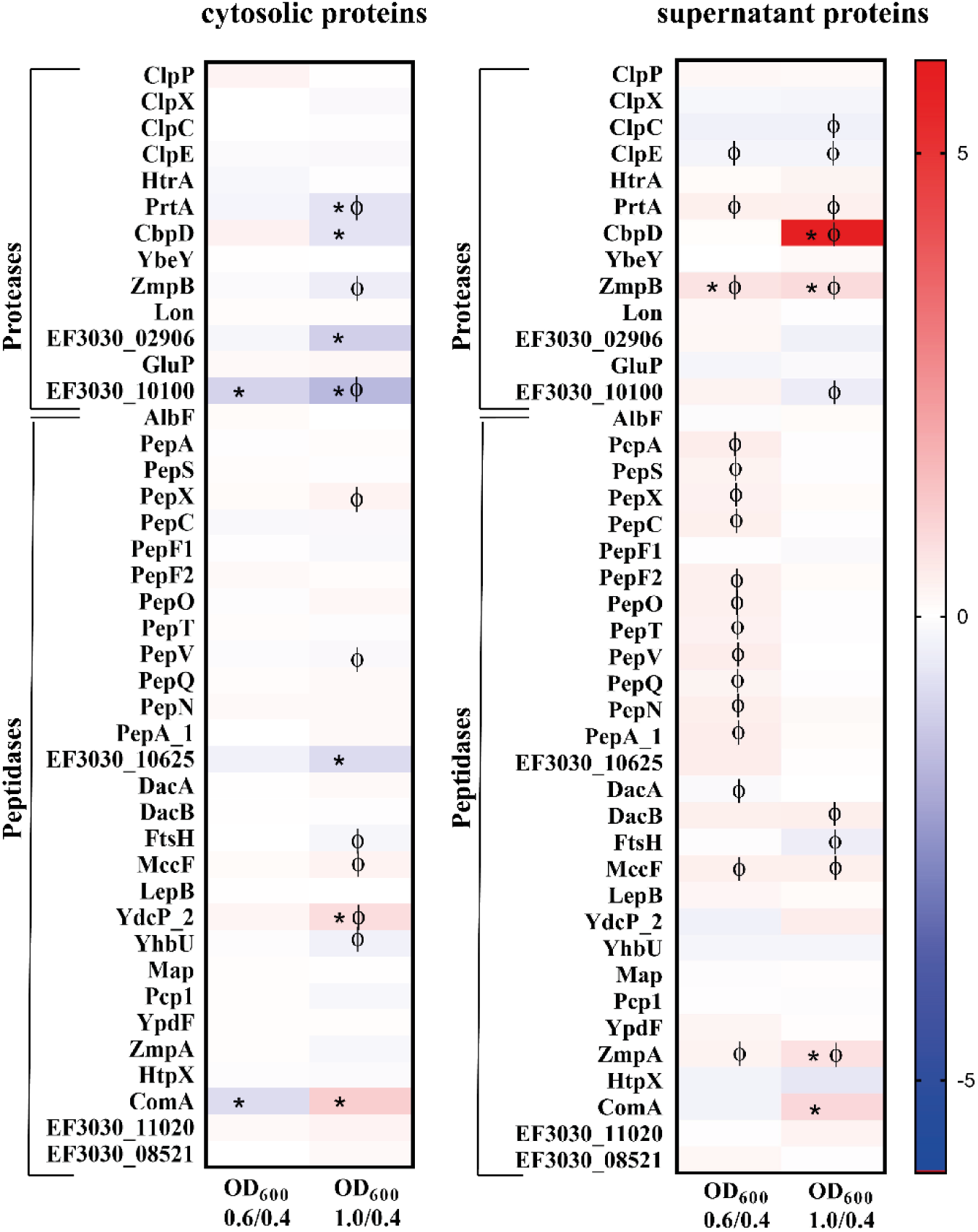
Heat maps illustrating a selection of proteases and peptidases with two or more peptides detected both in the cytosol and the supernatant of strain EF3030 by mass spectrometry analysis. The heat maps represent the signal log ratio values between mid-exponential phase to early exponential phase and late exponential phase to early exponential phase. Φ indicates significant values, p-value less than 0.05. * Indicates an absolute fold change greater than or equal to 1.5.

Among the pneumococcal peptidases, PepX and PepO exhibited increased abundance during late exponential phase in the cytoplasmic fraction compared to the early exponential phase, whereas PepS and AlbF showed a decrease in abundance during growth. Most of these peptidases had a decreased abundance during exponential growth in the extracellular fraction. AlbF was an exception, with a small increase in abundance observed at the late exponential phase.

## 4 Discussion

Understanding the dynamics of protein production during pneumococcal growth, colonisation, or invasive disease is essential for elucidating the molecular mechanisms underlying the pathophysiology of these pathobionts. This knowledge is important for identifying vaccine candidates and defining new targets for therapeutic interventions as well as developing sophisticated diagnostic tools. Our study employed an optimized minimal medium mimicking a nutrient-limited host compartment, such as the URT, to characterize the proteomic landscape of the colonising serotype 19F strain EF3030.

The nasopharyngeal environment is characterized by low CO2 levels, therefore we added bicarbonate included in our minimal medium, which provides an alternative source of hydrogen carbonate (HCO3^−^). This was particularly relevant given the findings of Burghout and colleagues, which highlighted the importance of carbonic anhydrase (PCA) activity in pneumococcal adaptation to CO2-poor conditions (Burghout et al., 2013). Carbonic anhydrase facilitates the reversible conversion of CO2 to bicarbonate, which is essential for fatty acid biosynthesis and polyglutamyl folate biosynthesis. No significant changes in carbonic anhydrase (MtcA1) levels were observed in our dataset, reinforcing the idea that it was not required due to the bicarbonate supplementation in the minimal medium. Additionally, several proteins from the *fab* gene cluster involved in fatty acid biosynthesis, such as FabD and FabG, showed significantly higher abundance in both the cytosolic fraction and the supernatant during the late exponential phase. Furthermore, the increased presence of FabK and FabZ in the supernatant during the late logarithmic phase suggests that the bacteria are directly utilizing bicarbonate to support growth and fatty acid synthesis. Because *S. pneumoniae* is auxotrophic for glutamine (Gln), we added this amino acid to RPMI 1640 (Härtel et al., 2011). Glutamine, taken up by a glutamine ABC transporter system, is crucial for pneumococcal metabolism and serves as a precursor for amino acids and nucleotides. Deprivation of glutamine impairs bacterial growth and virulence (Härtel et al., 2011).

Notably, both pneumococcal and streptococcal strains exhibited enhanced growth in our optimized minimal medium (CDM+), which was further supplemented with iron, manganese, and methionine. *S. pneumoniae* has a relatively low iron requirement, as it lacks a respiratory chain and does not contain cytochromes (Lanie et al., 2007). Nevertheless, the bacterium still requires iron for enzymes that contain iron-sulfur (FeS) clusters, such as anaerobic ribonucleotide reductase (Lanie et al., 2007). Previous studies have shown that *S. pneumoniae* strain TIGR4 exhibits reduced growth under iron-depleted conditions, which can be restored with iron supplementation (Gupta et al., 2009). In low-iron environments, pneumococci form long chains, indicative of impaired cell separation. Furthermore, mutations in the genes encoding the ABC transporters responsible for iron acquisition (PiaABCD and PiuBCDA) attenuated the virulence of *S. pneumoniae* in a mouse infection model (Brown et al., 2001; Brown et al., 2002).

Interestingly, in the present study, the strains, EF3030, TIGR4 and D39Δ*cps* exhibited clumping in CDM+, suggesting that the presence of iron may be a contributing factor. Previous studies also showed that elevated hemoglobin levels induce pneumococcal clustering and the secretion of factors that promote cell aggregation, suggesting adhesion and biofilm formation (Gupta et al., 2009).

Manganese is also an essential element in most organisms, serving as a cofactor for many enzymes involve in phosphorylation, hydrolysis, carbon metabolism, decarboxylation, and oxidative stress (Eijkelkamp et al., 2014). Pneumococcal proteins involved in manganese acquisition, such as MntE, NrdEF, SodD, PsaA, PsaR, and PhpP have a critical role in maintaining manganese homeostasis. When manganese levels are excessive, *S. pneumoniae* employs mechanism to cope the situation by reducing Mn uptake via the PsaABC transporter system, increasing Mn efflux (MntE), and inhibiting the Fe transport system (Martin et al., 2017). Our proteome data identified MntE, PsaA, MntR and PhpP in both cytosol and the supernatant. In addition to manganese, our minimal medium was also supplemented with uracil, which is a limiting factor for pneumococcal growth (Carvalho et al., 2013).

Building on this optimized medium, we focused our analysis on the protein profiles of *S. pneumoniae* EF3030 across three distinct growth phases: early (OD600 0.4), mid (OD600 0.6), and late exponential phase (OD600 1.0). Protein abundance was systematically assessed in both the cytoplasmic and culture supernatant fractions, providing a comprehensive view of the dynamic changes in protein abundance as the bacteria transitioned through different stages of growth. Notably, surface-associated and secreted proteins, which are crucial for pathogen-host interactions, exhibited significant variations in abundance depending on the growth phase. These proteins, including choline-binding proteins (CBPs), sortase-anchored proteins, and lipoproteins, are directly involved in bacterial adhesion, invasion, or immune evasion mechanisms and contribute to pneumococcal fitness. As such, they represent promising candidates for the development of vaccines and therapeutic strategies (Morsczeck et al., 2008; Rigden et al., 2003).

We identified over 1344 proteins that are common to both the supernatant and cellular fraction, suggesting a high rate of pneumococcal lysis in the medium already at the early stage of growth. This could be attributed to CbpD, a murein hydrolase, which showed a remarkable 54-fold increase in abundance in the extracellular fraction during the late exponential phase. However, even at lower levels early on, it may still trigger lysis in a subset of cells, potentially initiating a cascade effect.. CbpD causes lysis on its own and promotes an activation on LytA and LytC (Kausmally et al., 2005; Eldholm et al., 2009). The abundance of LytA increased in both the cytosol and the supernatant during exponential growth, although this increase was not statistically significant. In contrast, LytC was not detectable in our experimental conditions. We hypothesize that CbpD lyses non-competent sister cells, a process that is referred to as allolysis (Guiral et al., 2005) and the LytA from these cells promotes in-trans lysis of competent pneumococcal cells.

As *S. pneumoniae* colonises the nasopharynx, hydrogen peroxide (H₂O₂) production is thought to play a role in inhibiting competing microbes. SpxB, a pyruvate oxidase and the key enzyme for H₂O₂ production, was detected in higher abundance in the supernatant during the mid-exponential phase (OD600 0.6) and reduced during the later growth phase. The early expression of pyruvate oxidase during bacterial growth has also been reported in other studies (Pericone et al., 2000; Lee et al., 2006; Bättig and Mühlemann, 2008) coinciding with the time when pneumococci reach competence (Bättig and Mühlemann, 2008). Previous studies have demonstrated that the deletion of *spxB* abolishes spontaneous transformation, reduces the expression of early competence gene *comC*, and diminishes competence-associated DNA release (Bättig and Mühlemann, 2008). However, the role of SpxB products, H₂O₂ in these processes remain unclear. Interestingly, we failed to detect ComC, while we observed a higher abundance of ComB, a protein required for transformation, in the supernatant. Additionally, the late competence-associated proteins ComGA, which are essential for DNA uptake (Balaban et al., 2014), were found in significantly higher abundance both in the supernatant and cytosol, with a nearly 5-fold increase. Moreover, ComEA, a major component of DNA uptake apparatus for transformation was exclusively found in the supernatant, also increased in abundance during the late exponential phase. A study by Lui and colleagues have demonstrated that the receptor ComEA plays a significant role in the uptake of dsDNA during bacterial transformation, whereas HtrA, a serine protease, specifically degrades ComEA and ComEC to terminate the process (Liu et al., 2019). Other proteins involved in processing internalized single-stranded DNA (ssDNA), such as Dpr, DprA, and SsbB, were also found in higher abundance, although not always statistically significant, in both the cytosol and supernatant. *S. pneumoniae* cultivated in the minimal medium retains the ability to take up foreign DNA during the later stages of growth (Claverys et al., 2006). Altogether, these findings provide insights into the timing and regulation of competence and lysis in *S. pneumoniae*, highlighting the dynamic interplay between protein expression, autolysis, and DNA uptake during bacterial growth and colonisation. In addition, this demonstrates that the medium allows competence development in *S. pneumoniae* EF3030 during the exponential growth phase.

In our proteomic analysis of *S. pneumoniae* EF3030, we observed a significant increase in pneumolysin abundance (p-value < 0.05) from the early (OD600 0.4) to the late exponential phase (OD600 1.0) in the cytosolic fraction. However, this increase was not significant in the supernatant. In addition to pneumolysin, we identified several immunogenic proteins including PspC, PcpA, PrtA, PcsB, PhtD, PsaA, AliB, PavB, SatA, PnrA, PpmA, PspA, AmiA, LytN, and EF3030_RS10100 in both the cytosol and supernatant during the exponential growth phase. Serological profiling of pneumococcal proteins (He et al., 2024) and other studies have revealed that these proteins exhibit the highest IgG levels and are considered as vaccine candidates (Sempere et al., 2021; Khan and Pichichero, 2012; Tai, 2006). This suggests that the secretion of these proteins is a common phenomenon in *S. pneumoniae* during the exponential growth phase. Notably, both HtrA and PrtA, two serine proteases in *S. pneumoniae*, were detected in our dataset. While pneumococci typically harbour up to four secreted serine proteases (HtrA, PrtA, CbpG, and SFP) depending on the strain (Ali et al., 2021a), EF3030 lacks SFP. We did not detect CbpG in our experiments including tryptic peptides with up to two missed cleavages, however, we found both HtrA and PrtA in the early exponential phase (OD600 0.4). There was no significant fold change of HtrA during growth but PrtA showed a significant increase in the supernatant during exponential growth, further highlighting the dynamic protein secretion and during this phase.

The cultivation of EF3030 in the minimal medium also revealed a higher abundance of several zinc-acquisitionproteins, including the AdcA and AdcAII transporters, the repressor AdcR, and the pneumococcal histidine triad (Pht) proteins. The Pht proteins, in particular, are known virulence factors in *S. pneumoniae*, with promising potential in preclinical vaccine trials aimed at preventing pneumococcal colonization and disease (Adamou et al., 2001; Godfroid et al., 2011; Denoël et al., 2011; Seiberling et al., 2012; Plumptre et al., 2013). Studies have shown that the genes encoding for AdcAII and PhtD are part of an operon regulated by the repressor AdcR. PhtD serves as an extracellular Zn²⁺ scavenger, acting as a metallophore for zinc acquisition and potentially transferring captured Zn²⁺ to the AdcAII transporter or forming a complex with AdcAII, linking zinc sensing to virulence(Loisel et al., 2011) In the presence of zinc, AdcR directly represses the *adcRCBA* genes and *adcII, phtA, phtB, phtD*, and*phtE* (Shafeeq et al., 2011).

Studies have shown that a lack of zinc (Zn) increases the risk of severe invasive pneumococcal infections and mortality. Research by (Strand et al., 2001) provides direct evidence that individuals with Zn deficiency experience more severe pneumococcal infections. Furthermore, a study by (Strand et al., 2003) reported that Zn deficiency not only worsens the infection but also weakens the body’s ability to produce specific antibodies against PspA. This leads to higher bacterial colonization in mucosal tissues, increasing the likelihood of invasive disease and death. On the other hand, research by (Bhandari et al., 2002) demonstrated that providing zinc supplements to children in developing countries significantly lowers the occurrence of pneumonia.

Given that our experiments were conducted under zinc-limiting conditions, the significant increase in Zn²⁺ transporters and Pht proteins is consistent with the bacterium’s need to scavenge zinc from its environment. This aligns with previous studies showing that Zn depletion weakens host immunity and increases pneumococcal colonization and virulence, further emphasizing the role of Zn in controlling pneumococcal infections. These findings suggest that supplementing the culture medium with zinc could reduce or eliminate the high abundance of these proteins, highlighting the adaptive mechanisms *S. pneumoniae* employs to thrive in zinc-limited conditions during infection.

Finally, a comprehensive understanding of the proteome profile during *S. pneumoniae* growth provides valuable insights into the molecular mechanisms underlying its colonisation and pathogenicity. This knowledge can be leveraged to identify potential vaccine targets by highlighting key proteins involved in host interaction, immune evasion, and virulence. Additionally, it could aid in the development of strategies to block critical resistance factors, such as antibiotic efflux pumps, or surface adhesins, thereby enhancing therapeutic options. Ultimately, these findings contribute to a more detailed understanding of pneumococcal biology, which is essential for designing more effective treatments and preventive measures against this pathogen.

## 5 Conclusion

This study provides a detailed proteomic profile of *S. pneumoniae* EF3030, revealing growth-dependent changes in both the cellular and extracellular proteomes. The identified proteins, particularly those involved in transport, regulation, metabolism, and virulence, offer valuable insights into the bacterial adaptation to varying growth conditions and the potential targets for therapeutic interventions. The characterization of these proteomic shifts significantly advances our understanding of pneumococcal physiology and its response to host environments.

## 7 Author Contributions

SD: Investigation, Data curation, Methodology, Formal Analysis, Visualization, Writing-original draft; LB: Software, Formal Analysis, Validation, Visualization, Writing-review & editing; GB: Formal Analysis, Supervision, Writing-review & editing; MGS: Software, Formal Analysis, Writing-review & editing; RS: Investigation, Methodology, Resources, Visualization, Writing-review & editing; LS: Investigation, Software, Data curation, Formal Analysis, Validation, Writing-review & editing; UV: Supervision, Validation, Project administration, Funding acquisition, Resources, Writing-review & editing; SH: Conceptualization, Supervision, Formal Analysis, Project administration, Validation, Funding acquisition, Resources, Writing-review & editing. All authors contributed to the article and approved the submitted version.

## 9 Acknowledgments

We like to thank Katrin Schoknecht (University Medicine Greifswald), Birgit Rietow (University of Greifswald) and Stefan Bock (University of Greifswald) for technical assistance, Vera Fuentes Moreno and Luca Cagnini for supporting experimental work, Prof. Dr. Anders Hakansson, Lund University, Sweden for kindly providing the 19F, EF3030 strain. The data illustrations were created with GraphPad Prism, version 8.4.3 (686), Microsoft Excel and R version 4.4.1. The language was improved using OpenAI, ChatGPT.

## 9 Data Availability Statement

The datasets [GENERATED/ANALYZED] for this study can be found in the [NAME OF REPOSITORY] [LINK]. Please see the “Availability of data” section of [Materials and data policies in the Author guidelines](https://www.frontiersin.org/guidelines/policies-and-publication-ethics#materials-and-data-policies) for more details.

Reviewer Access via the PRIDE website (https://www.ebi.ac.uk/pride/) using the following details

*Project accession: PXD062379*

*Token: aIkOo8bG7wCK*

## Supplementary Material

### 1 Supplementary Figures

**Supplementary Figure S1:**
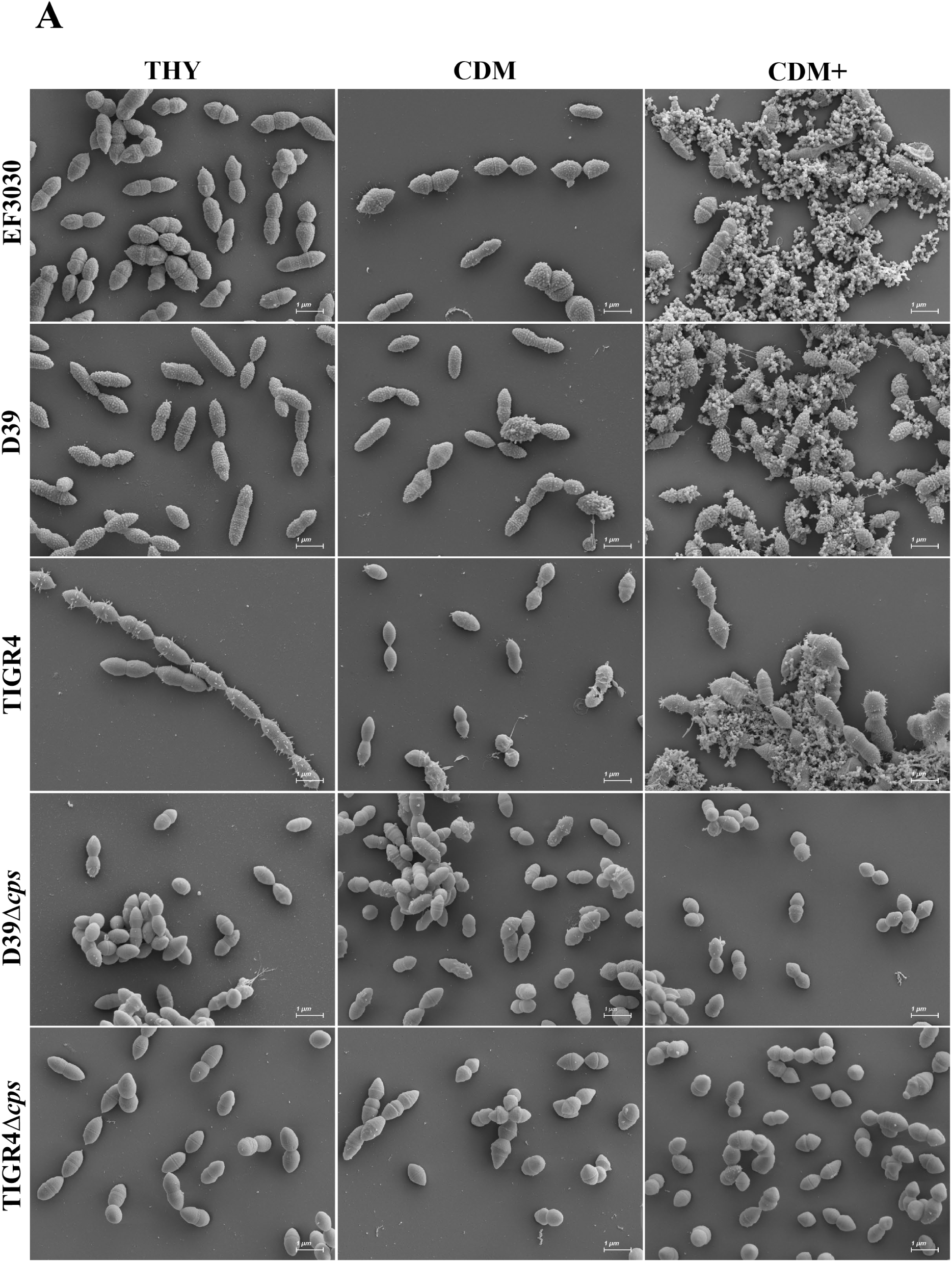
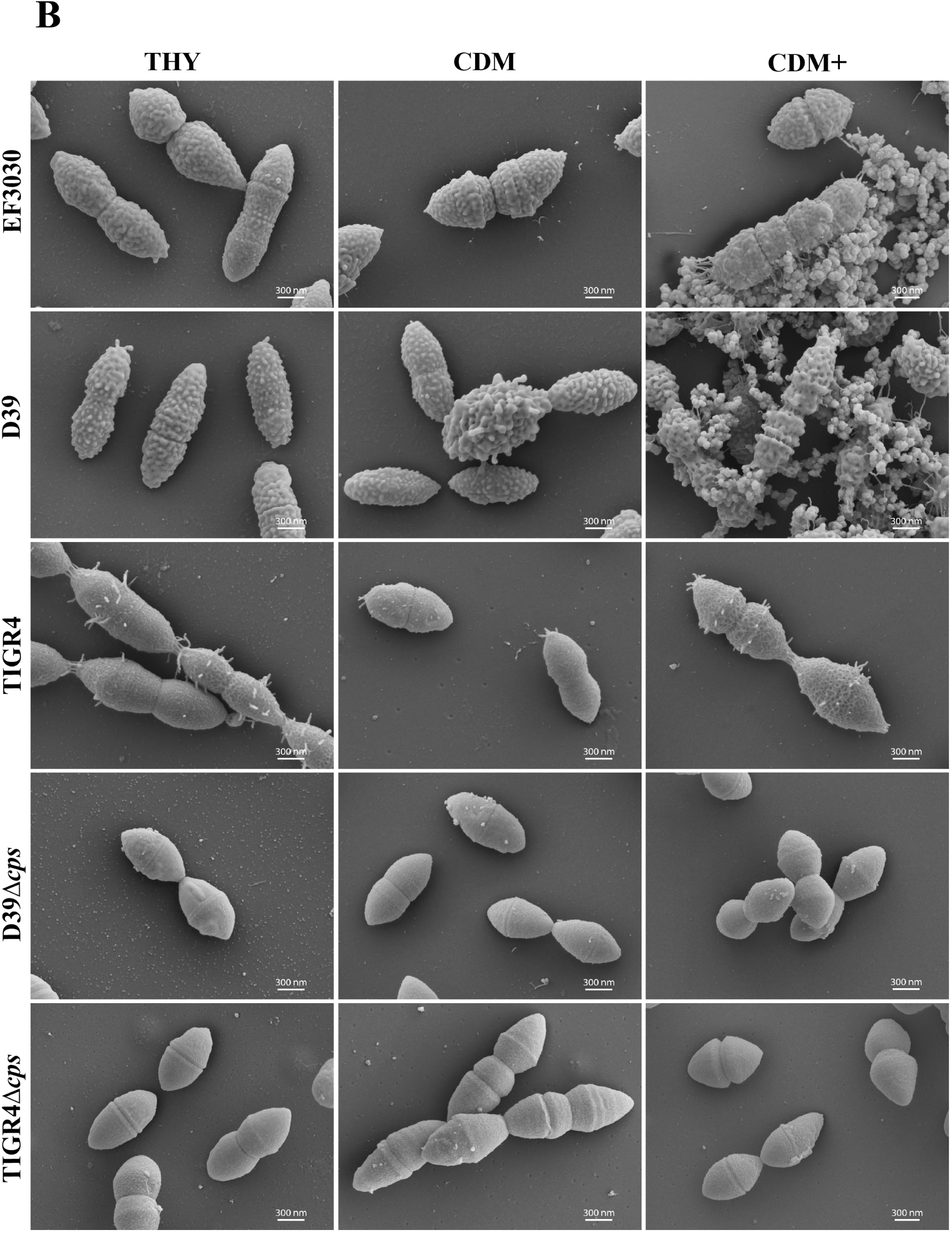
Scanning electron micrographs of *S. pneumoniae* cultivated in THY, CDM, or CDM**+.** The micrographs were taken at a magnification of 10,000x, scale bars = 1 µm (A) and at a higher magnification of 30,000x, scales bar = 300 nm (B).

**Figure S2:**
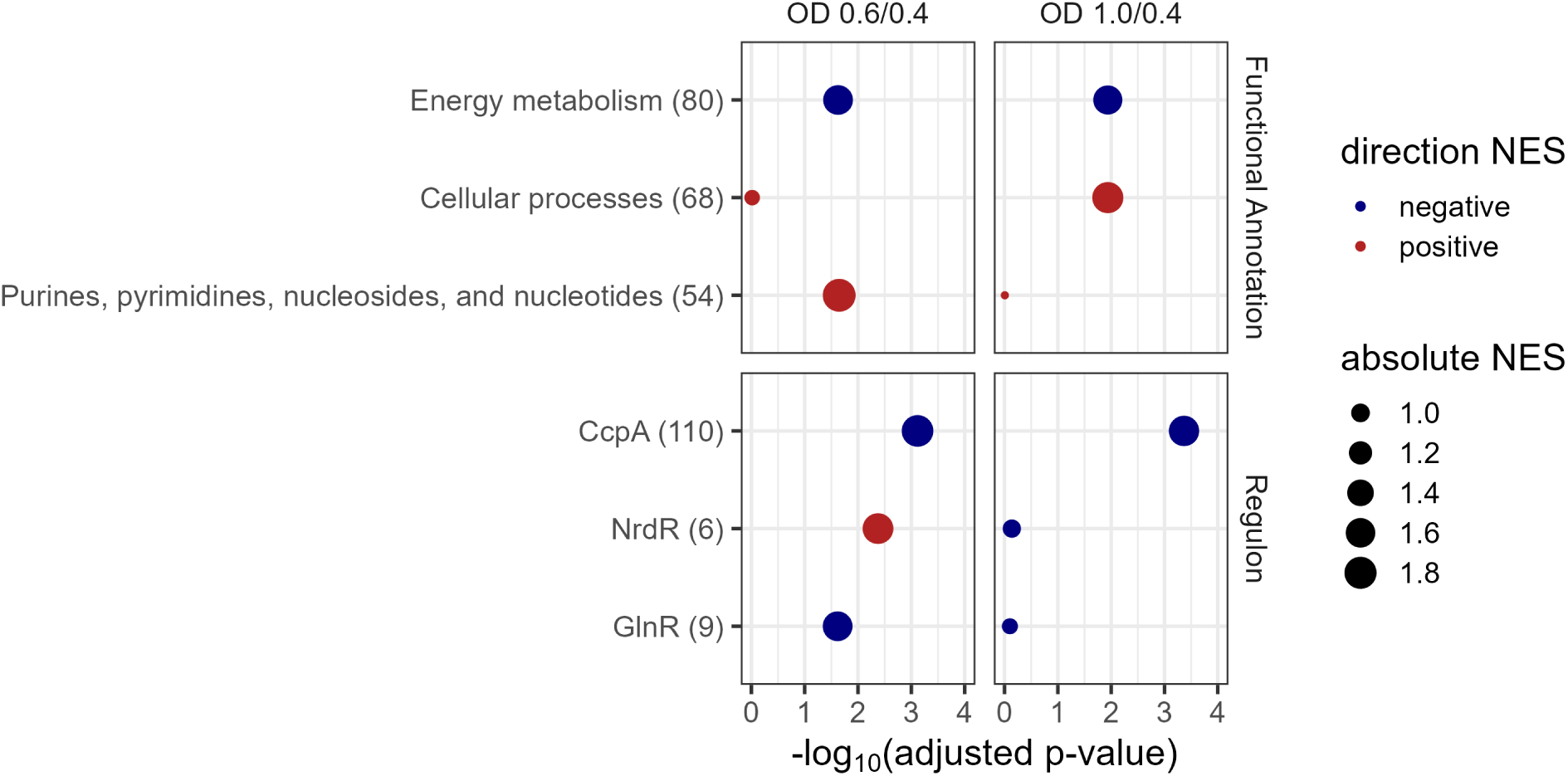
Gene set enrichment analyses (GSEA) comparing different biological functions and regulatory networks between conditions. X-axis represents log_10_ (adjusted p-value), y-axis shows Functional annotation (top panel) and Regulon (bottom panel). Each dot represents a pathway or regulon indicated by “Red: positively enriched” and “Blue: negatively enriched”. The size of the dots reflects the absolute Normalized Enrichment Score (NES) indicating magnitude of enrichment.

**Supplementary Figure S3:**
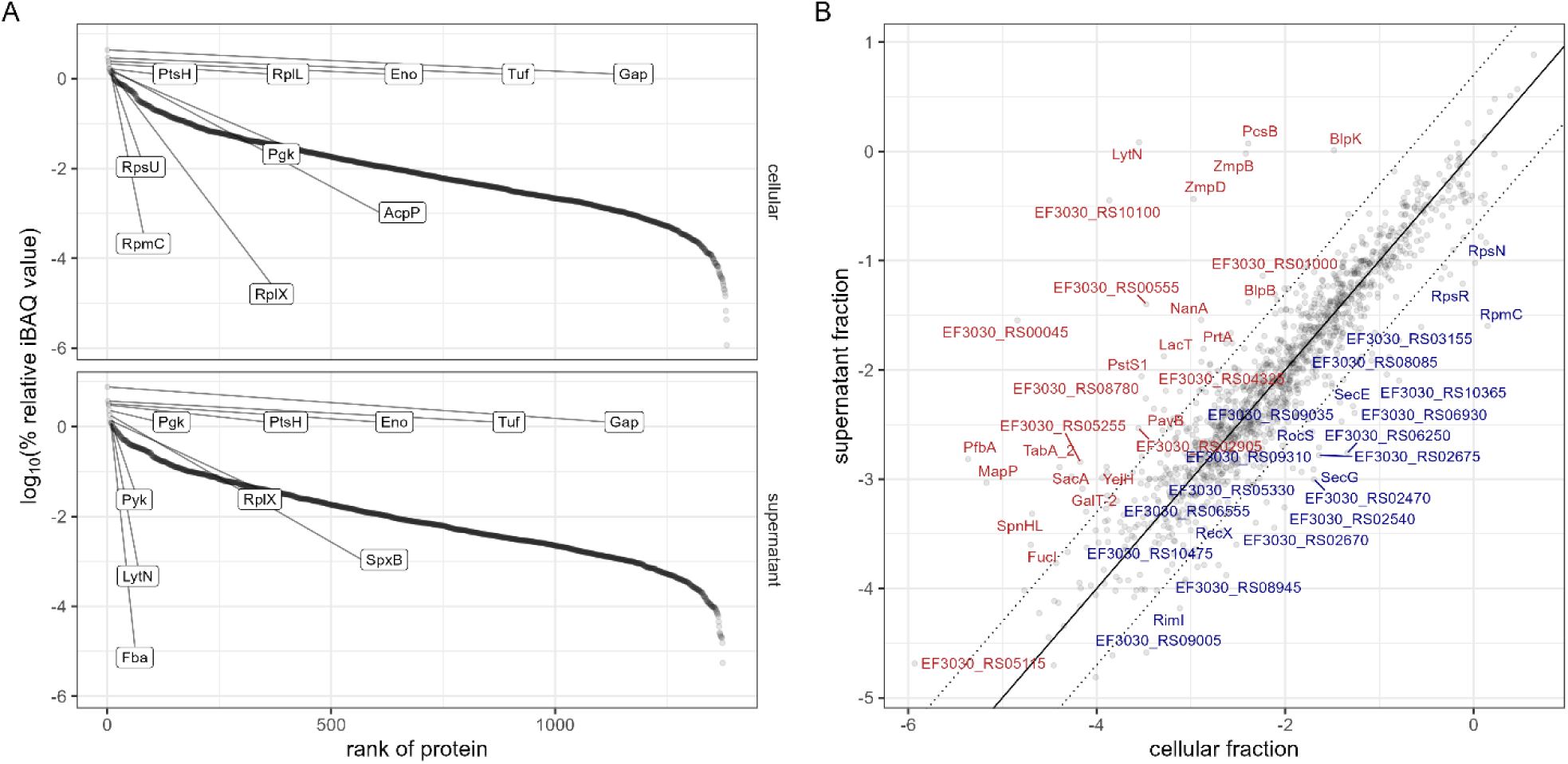
Protein composition of the cellular and supernatant fraction of the proteome at OD_600_ 1.0. (A) Display of the range of detected abundances scaled as relative iBAQ values in the cellular and supernatant fraction. The top 10 most abundant proteins are labelled. (B) Comparison of the log_10_-scaled relative iBAQ value per protein in the cellular and supernatant fraction. Solid line displays perfect correlation; dotted lines display 5-fold difference of relative abundance between both fractions. Proteins differing at least 10-fold are labelled. Proteins in higher relative abundance in the supernatant are coloured red and proteins in higher relative abundance in the cellular fraction are coloured blue.

### 2 Supplementary Tables

**Table S1:** Pneumococcal and streptococcal strains used for studying the growth in the modified minimal medium (CDM+)

| Species and strain | Relevant characteristics | Reference |
| --- | --- | --- |
| *S. pneumoniae* |  |  |
| D39 | wild-type strain, serotype 2 | NCTC 7466 |
| D39Δ*cps* | Allele exchange mutant *cps::aphA3*, capsule deficient, Km<sup>r</sup> | (Rennemeier et al. 2007) |
| TIGR4 | wild-type strain, serotype 4 | (Tettelin et al. 2001) |
| TIGR4Δ*cps* | Allele exchange mutant *cps::aphA3*, capsule deficient, Km<sup>r</sup> | (Schulz et al. 2014) |
| EF3030 | wild-type strain, serotype 19F | (Junges et al. 2019) |
| Invasive clinical isolate | 8 | RLS* (RWTH-Aachen) |
| Invasive clinical isolate | 12F | RLS (RWTH-Aachen) |
| Invasive clinical isolate | 22F | RLS (RWTH-Aachen) |
| *S. mutans* | UA159 | ATCC |
| *S. suis* | Strain 10, serotype 2 | (Baums et al. 2009) |
| *S. pyogenes* | Group A Streptococcus 5448 | (Kaul et al. 1997) |
| *S. agalactiae* | *S. agalactiae*, GBS STSS/NF-HH NSTI, STSS; Serotype Ib; ST8; Stockholm (Sweden) | (Sendi et al. 2009) |

Km<sup>r</sup>, kanamycin resistant; \*RLS: Reference Laboratory for Streptococci

**Table S2:** Specific information of the Reversed phase liquid chromatography (RPLC)

|  |  |
| --- | --- |
| *instrument* | Ultimate 3000 RSLC (Thermo Scientific) |
| *trap column* | 75 µm inner diameter, packed with 3 µm C18 particles (Acclaim PepMap100, Thermo Scientific) |
| *analytical column* | Accucore 150-C18, (Thermo Fisher Scientific)<br>25 cm x 75 µm, 2,6 µm C18 particles, 150 Å pore size |
| *buffer system* | binary buffer system consisting of 0.1% acetic acid in HPLC-grade water (solvent A) and 100% ACN in 0.1% acetic acid (solvent B) |
| *flow rate* | 300 nl/min |
| *gradient* | 0min-2% B<br>2min-5% B<br>10min-7% B<br>70min-25% B<br>75min-40% B<br>77min-90% B<br>83min-90% B<br>85min-2% B<br>95min-2%B |
| *column oven temperature* | 40°C |

**Table S3:** Specific information of the Mass-spectrometry analysis in the data-independent acquisition mode.

|  |  |
| --- | --- |
| *instrument* | Orbitrap Exploris™ 480 |
| *electrospray* | Nanospray Flex™ Ion Source |
| *operation mode* | data-independent |
| **Full Scan Properties** |  |
| *MS scan resolution* | 120000 |
| *AGC target* | 3e6 (300%) |
| *maximum ion injection time for the MS scan* | 60 ms |
| *Scan range* | 350 to 1200 m/z |
| *Microscans* | 1 |
| *Polarity* | positive |
| *RF Lens* | 50% |
| *Spectra data type* | profile |
| **DIA Properties (MS2)** |  |
| *Resolution* | 30,000 |
| *maximum ion injection time for the MS/MS scans* | auto |
| *Normalized AGC target* | 3E6 |
| *Spectra data type* | profile |
| *Microscans* | 1 |
| *isolation window* | 66 |
| *Isolation window width* | 13 m/z |
| *Window overlay* | 2 m/z |
| *Fixed first mass* | 200 |
| *HCD collision energy* | 30% |

**Table S4:**
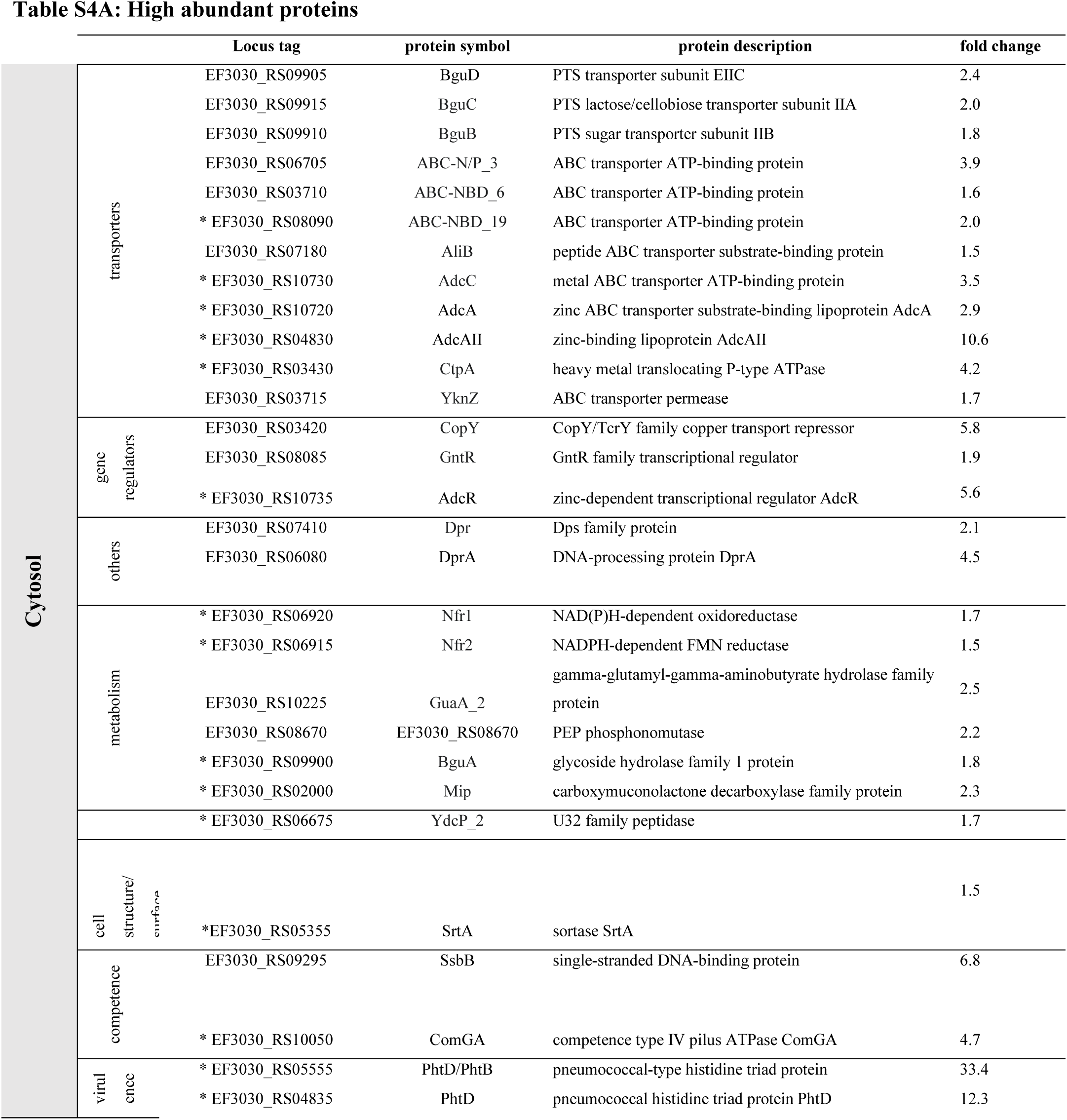
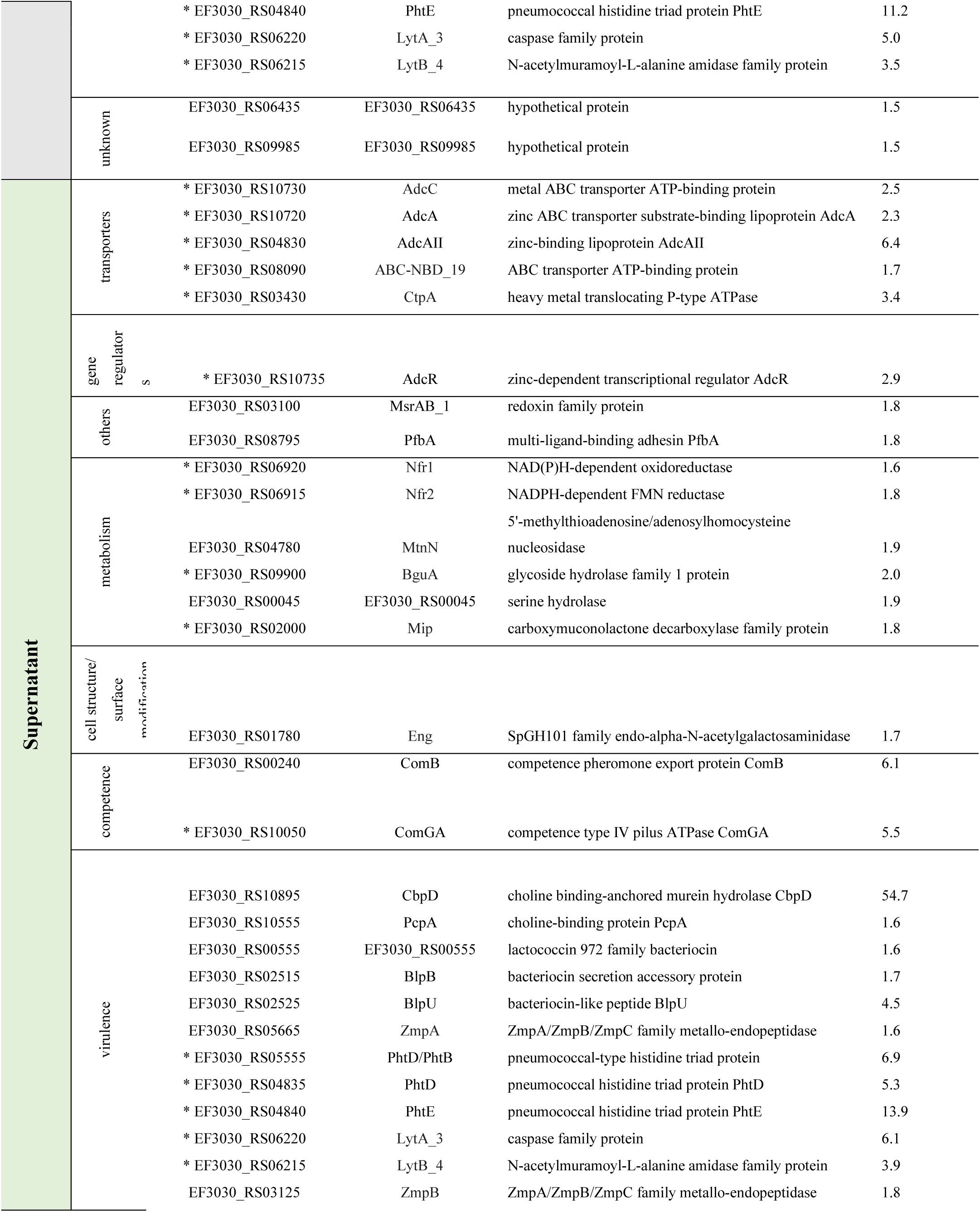
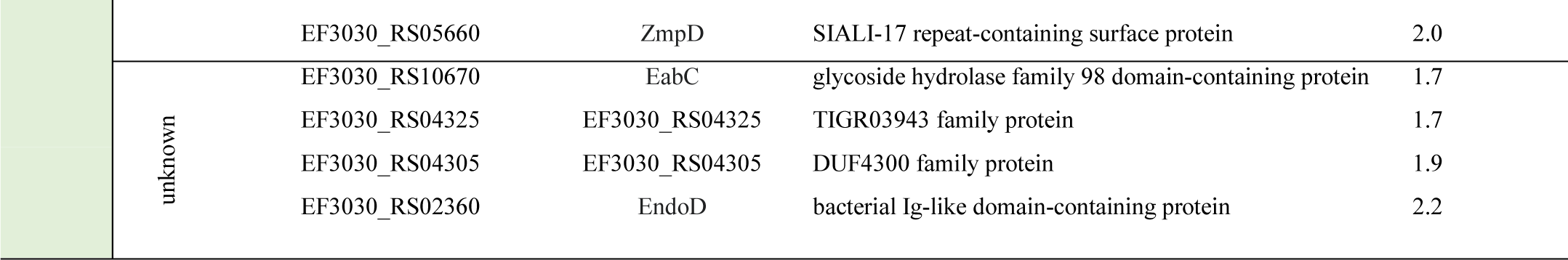
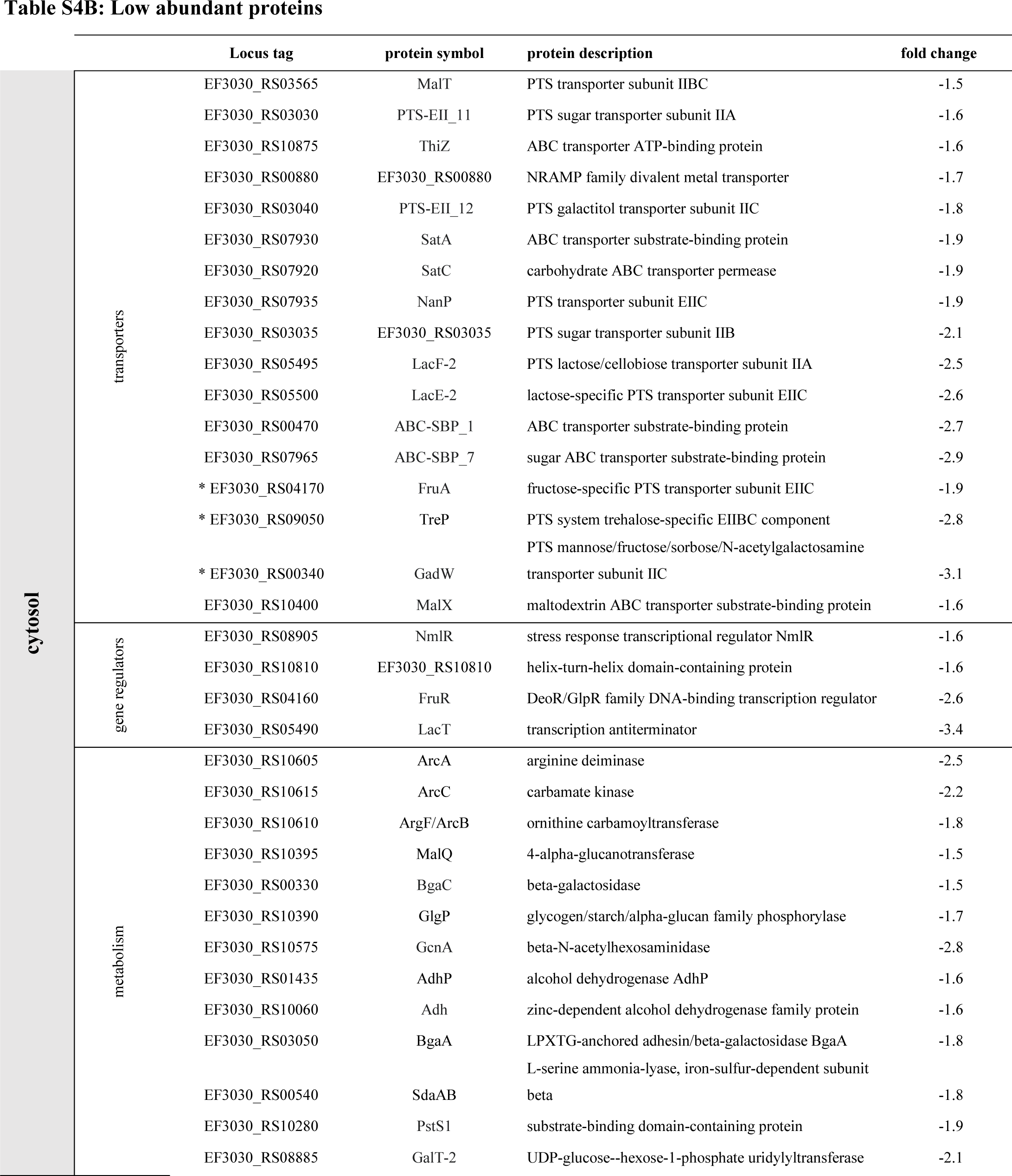
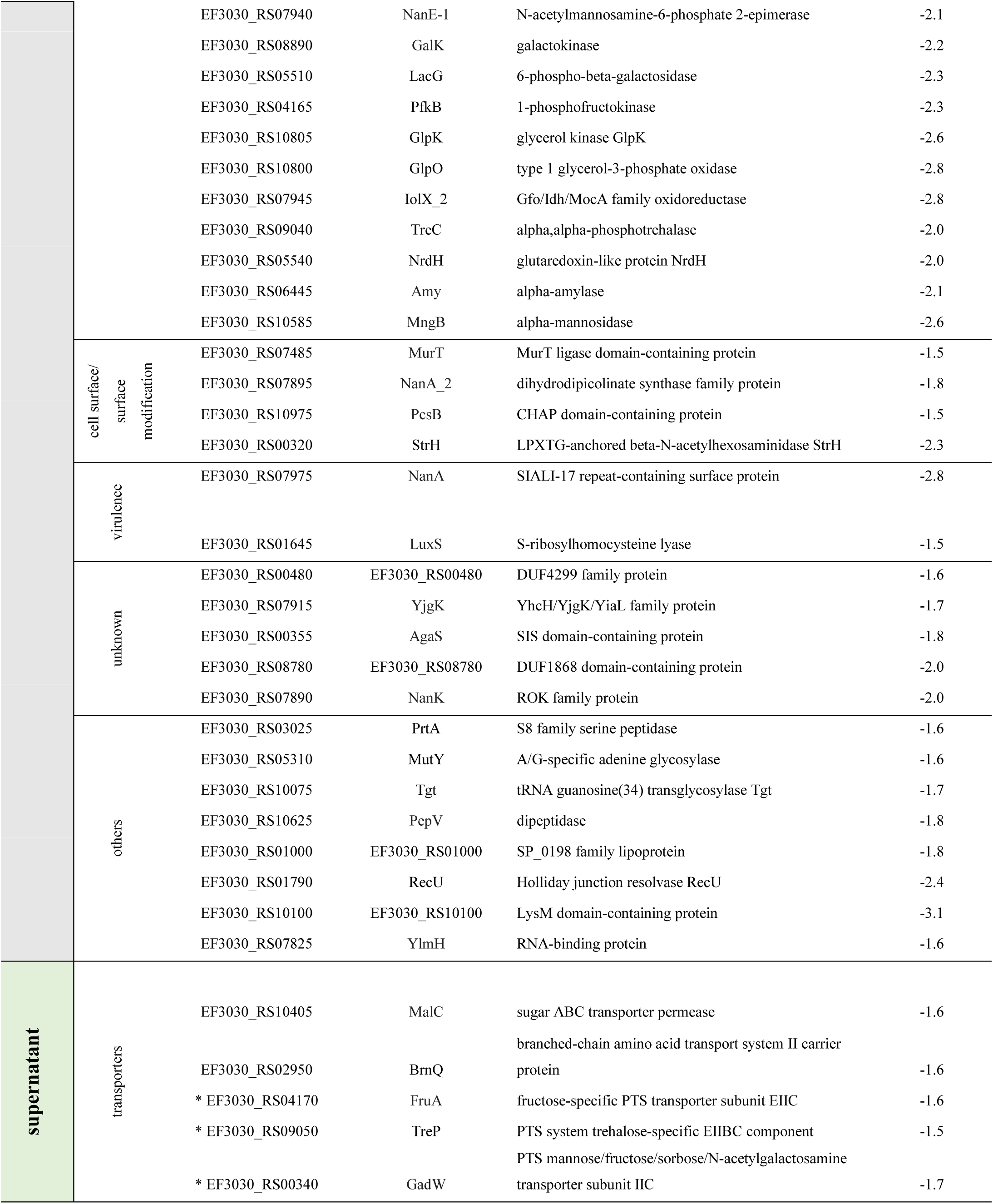
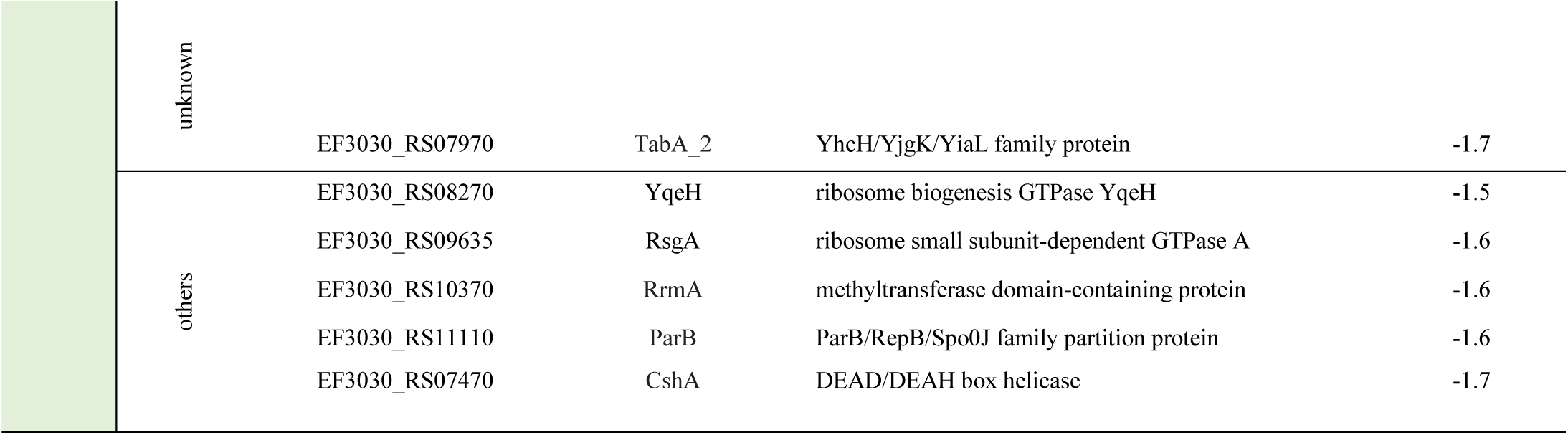
Proteins with significant changes in abundancies during exponential growth. The proteins identified in the cytosolic fraction and the supernatant fraction with a significant change in abundance during the late exponential growth (OD_600nm_ 1.0) compared to the early exponential growth (OD_600nm_ 0.4). The high abundant proteins and low abundant proteins detected in both the cytosolic and supernatant fractions are marked by “*”.

